# Integrative analysis of metabolite GWAS illuminates the molecular basis of pleiotropy and genetic correlation

**DOI:** 10.1101/2022.04.02.486791

**Authors:** Courtney J. Smith, Nasa Sinnott-Armstrong, Anna Cichońska, Heli Julkunen, Eric Fauman, Peter Würtz, Jonathan K. Pritchard

**Author notes:** Corresponding Authors: Courtney J. Smith, Nasa Sinnott-Armstrong, Jonathan K. Pritchard.

## Abstract

Pleiotropy and genetic correlation are widespread features in GWAS, but they are often difficult to interpret at the molecular level. Here, we perform GWAS of 16 metabolites clustered at the intersection of amino acid catabolism, glycolysis, and ketone body metabolism in a subset of UK Biobank. We utilize the well-documented biochemistry jointly impacting these metabolites to analyze pleiotropic effects in the context of their pathways. Among the 213 lead GWAS hits, we find a strong enrichment for genes encoding pathway-relevant enzymes and transporters. We demonstrate that the effect directions of variants acting on biology between metabolite pairs often contrast with those of upstream or downstream variants as well as the polygenic background. Thus, we find that these outlier variants often reflect biology local to the traits. Finally, we explore the implications for interpreting disease GWAS, underscoring the potential of unifying biochemistry with dense metabolomics data to understand the molecular basis of pleiotropy in complex traits and diseases.

## Introduction

A central challenge in the field of human genetics is understanding the mechanism of how genetic variants influence complex traits and diseases. Genome-wide association studies (GWAS) have begun characterizing the genetic architecture of complex traits, but the molecular mechanisms connecting genetic variants to these traits are rarely understood. This is particularly true for understanding pleiotropy, when a variant affects multiple traits [1]. It is possible to estimate the genetic correlation between traits [2, 3], but it is often unclear what contributes to this at a molecular or physiological level. A handful of *in vitro* disease-focused “post-GWAS” studies have convincingly shown the mechanisms driving pleiotropy of individual key associations [4, 5]; however, these studies are highly specific and time-consuming. Developing statistical and computational approaches to identify putative molecular mechanisms is invaluable to advancing our understanding of where and how pleiotropic GWAS variants act.

In this study, we use metabolites as model traits to understand pleiotropic features of genetic architecture. Metabolites are small molecules interconverted by a series of biochemical pathways, and are an appealing model system for studying pleiotropy because their pathways are typically well-documented and biologically simpler than those underlying other complex traits [6, 7]. Previous work in Mendelian genetics has identified inborn errors of metabolism (IEM) in many enzymes [8]. Metabolite GWAS, which have long observed pervasive pleiotropy at these IEM genes and other loci [9, 10], offer a potential opportunity to further explore the relationships between intermediate molecules and disease outcomes at scale. Here, we jointly analyzed GWAS results of 16 plasma metabolites from the Nightingale Health Nuclear Magnetic Resonance (NMR) Spectroscopy platform in nearly 100,000 individuals in the UK Biobank [11] (Figure 1; see Methods). These 16 metabolites included glucose, pyruvate, lactate, citrate, isoleucine, leucine, valine, alanine, phenylalanine, tyrosine, glutamine, histidine, glycine, acetoacetate, acetone, and 3-hydroxybutyrate. They were chosen based on their biochemical proximity to each other, their relevance to health and disease, and because the genes and enzymes involved in their metabolism are well-characterized. They play especially important roles in energy generation and energy storage pathways such as glycolysis, the citric acid cycle, amino acid metabolism and ketone body formation. They are relevant to many metabolic diseases including type 2 diabetes [12, 13, 14], cardiovascular disease [15], and non-alcoholic fatty liver disease [16].

**Figure 1:**
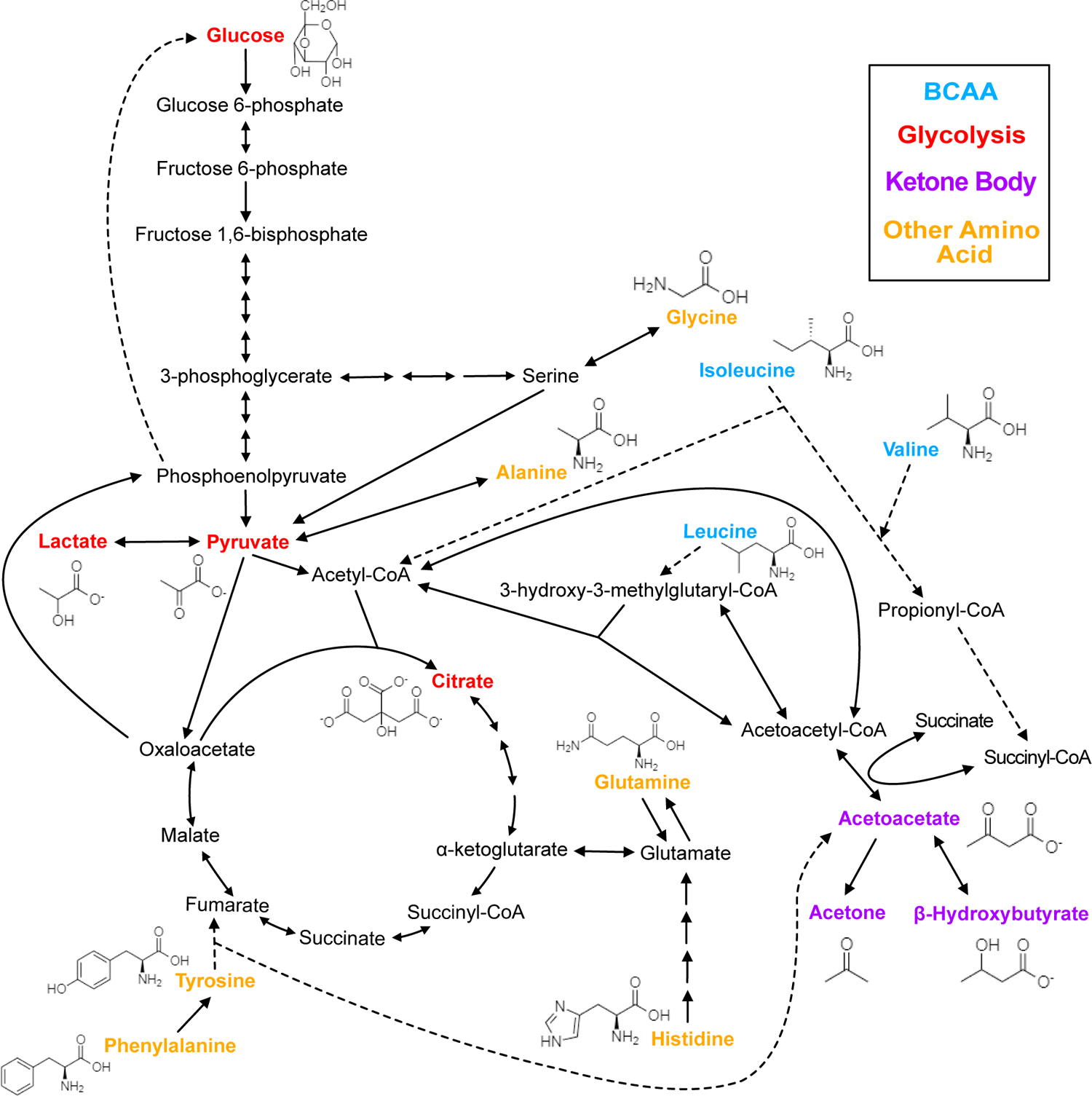
Biochemistry of relevant metabolites. Pathway diagram and molecular structure of relevant metabolites, colored by their biochemical groups. The pathway diagram was curated from multiple resources (see Methods). All solid lines represent a single chemical reaction step. Dotted lines represent a simplification of multiple steps. For simplicity, only a subset of all the reactions each metabolite participates in is shown. Genes encoding the enzymes that catalyze the above chemical reactions are known and presented in Supplementary Figure S4.

Numerous GWAS have begun characterizing the genetic architecture of metabolites and found them to be heritable and polygenic [17, 18]. Recent metabolite studies have shown that leveraging information about the biochemical pathways relevant to a given metabolite [19, 20, 21, 7] can allow for more interpretable gene annotation of GWAS hits. This has led to the dissection of individual associations of biomarkers, such as lipids [22], glycine [23], and intermediate clinical measures [24], with cardiometabolic and other diseases. The pervasive pleiotropy at these GWAS loci with other metabolites as well as disease [24, 25] suggests the potential of utilizing these data for investigating the mechanism of pleiotropic effects as a core component of genetic architecture. While recent GWAS have begun jointly investigating multiple metabolites [26, 27, 28], they have yet to do so in the context of their biochemical pathways.

In this paper, we demonstrate that investigating the effects of pleiotropic variants on biologically-related metabolites allows for a better understanding of why these variants have their observed joint effects. Our results reveal striking heterogeneity in genetic correlation across the genome and provide a biologically intuitive basis for understanding this heterogeneity. Together, this allows us to dissect the molecular basis of metabolic disease GWAS variants and enables us to directly define the mechanism relating an example variant to its associated disease.

## Results

### Insights into the shared genetic architecture of biologically-related metabolites

We chose 16 metabolites from the 249 available through the Nightingale NMR platform in a subset of the UK Biobank (Figure 1; see Methods). These 16 metabolites were selected based on their biochemical proximity, relevance to health and disease, and because the genes and enzymes involved in their metabolism are well-characterized. We classified the 16 metabolites into four groups based on shared biochemistry: Glycolysis (glucose, pyruvate, lactate, citrate), Branched Chain Amino Acid (BCAA; isoleucine, leucine, valine), Other Amino Acid (alanine, phenylalanine, tyrosine, glutamine, histidine, glycine), and Ketone Body (acetoacetate, acetone, 3-hydroxybutyrate). Trait measurements were log-transformed and adjusted for relevant technical covariates. After outlier removal, we obtained a primary dataset of 94,464 genotyped European-ancestry individuals with data for all 16 metabolites. Additionally, we performed an ancestry-inclusive GWAS of all 98,189 individuals with complete metabolite data for followup analysis.

We first sought to characterize the genetic architecture underlying these metabolites by performing GWAS for each (Supplementary Figure S1). Hits from individual GWAS were clumped with an *r*^2^ of 0.01 per megabase, combined across metabolites, then pruned to the SNP with the most significant P-value within 0.1 cM. This resulted in 213 lead variants with a genome-wide significant association in at least one metabolite, referred to as the metabolite GWAS hits. Glycine had the largest number of significant associations with 77 hits (Figure 2a). There were 47 variants with significant associations in more than one metabolite, including rs2939302 (near the gene *GLS2*) which was significant in 9 of the 16 metabolites, and rs1260326 (*GCKR*) which was significant in 8. Glycine also had the highest total SNP heritability of 0.284 (Supplementary Table S1 and Supplementary Figure S2).

**Figure 2:**
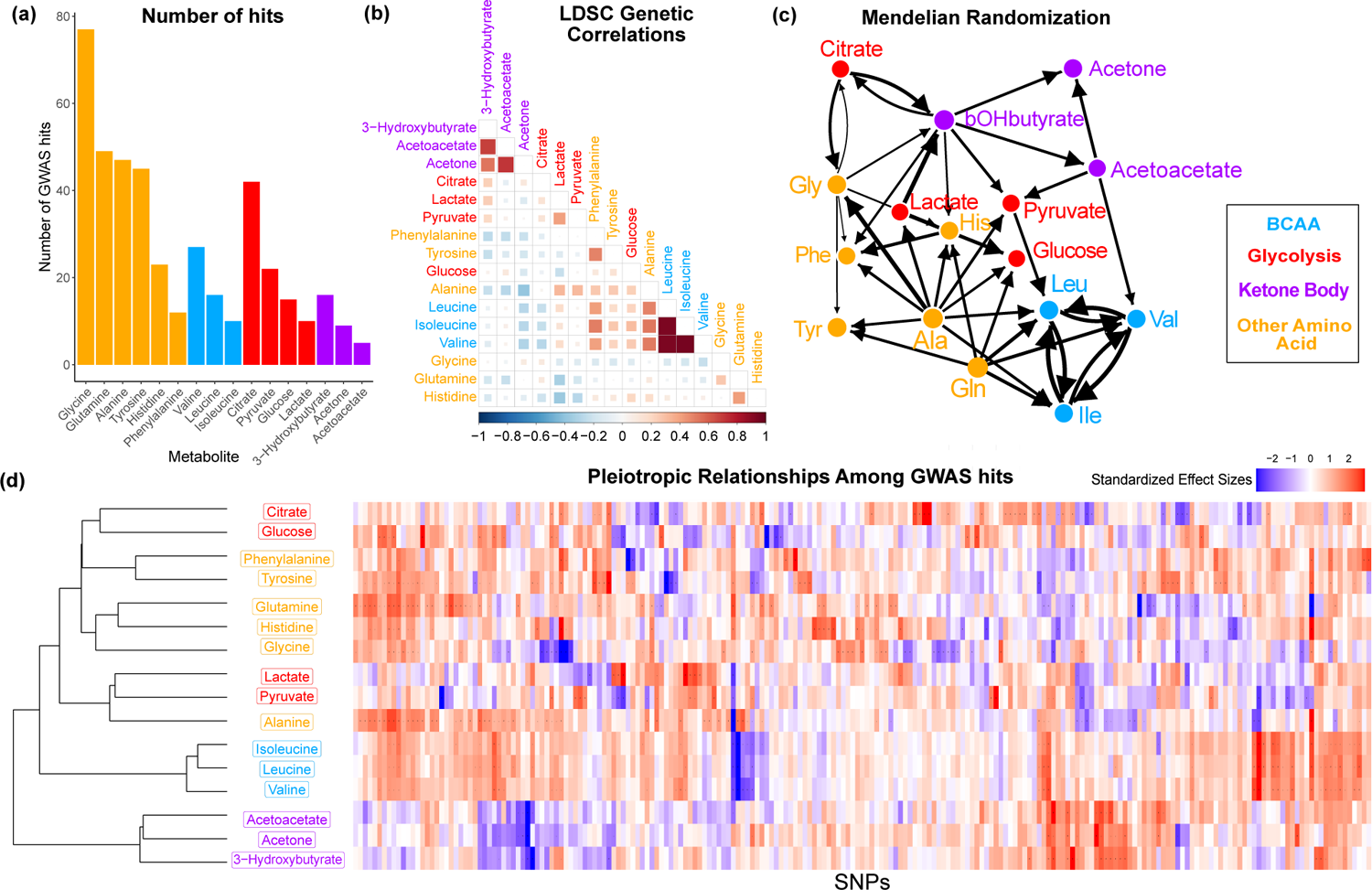
Overview genetic architecture of metabolites. **a.** Number of GWAS hits per metabolite. **b.** Pairwise LDSC genetic correlations between the metabolites, clustered by genetic correlation. **c.** Mendelian Randomization weighted results between the metabolites. **d.** Biclustered standardized effect size in each metabolite for the 213 metabolite GWAS hits. For visualization, effect sizes were divided by the standard error then inverse normal transformed and standardized. Each variant was aligned to have a positive median score across metabolites.

To understand the shared genetics of these metabolites, we then investigated the extent of pleiotropy between and within biochemical groups. In order to examine this, we first calculated pairwise LDSC genetic correlation across the 16 metabolites. We found substantial genome-wide sharing for many pairs of metabolites, especially for metabolites within the same biochemical group (Figure 2b; phenotypic correlation in Supplementary Figure S3). We then explored pleiotropic effects beyond the polygenic background by examining the structure within the metabolite GWAS hits. Pairwise Mendelian Randomization (MR) between the metabolites emphasized the intertwined nature of these traits (Figure 2c). Despite only taking into account genetic effects, MR largely clustered metabolites in a way that reflects their biochemical groups. The extensive pleiotropy across the 16 traits, with similar sharing inside biochemical groups, is also illustrated by the structure visible in the normalized effect sizes for each metabolite GWAS hit (Figure 2d). Together, these analyses support substantial, but not always consistent, genetic overlap between the traits, particularly in the polygenic components. In the remainder of this paper, we will seek a deeper understanding of the biochemical relationships between genotypes and metabolite levels.

### Characterizing the biological functions of candidate genes

An important step in understanding the pathway level mechanisms of variants is knowing which gene a variant is affecting and how that gene relates to the biology of the pathway. Different types of genes influence trait biology through distinct mechanisms. Metabolite biology is documented in genetic and biochemical databases based on the extensive history of biochemical research (Supplementary Table S2). Thus, we developed a pipeline for annotating the 213 metabolite GWAS hits with a single most likely gene using gene proximity and manual curation of these databases (Supplementary Table S3 and Supplementary Figure S5; see Methods). We annotated 68 variants with genes encoding pathway-relevant enzymes (25-fold enrichment, Poisson rate test P < 2e-16), 46 with genes encoding transporters (5.2-fold enrichment, P = 9e-16), and 30 with genes encoding transcription factors (7-fold enrichment among liver marker TFs, P = 3e-5; Figure 3). Overall, 69% of variants were assigned to the closest gene and 49% of variants assigned to a pathway-relevant enzyme gene were assigned known inborn errors of metabolism (IEM) genes [8]. The substantial enrichment for biologically interpretable variants suggests that examining the genetic basis of these traits will allow for the development of hypotheses around relevant molecular mechanisms underlying pleiotropy.

**Figure 3:**
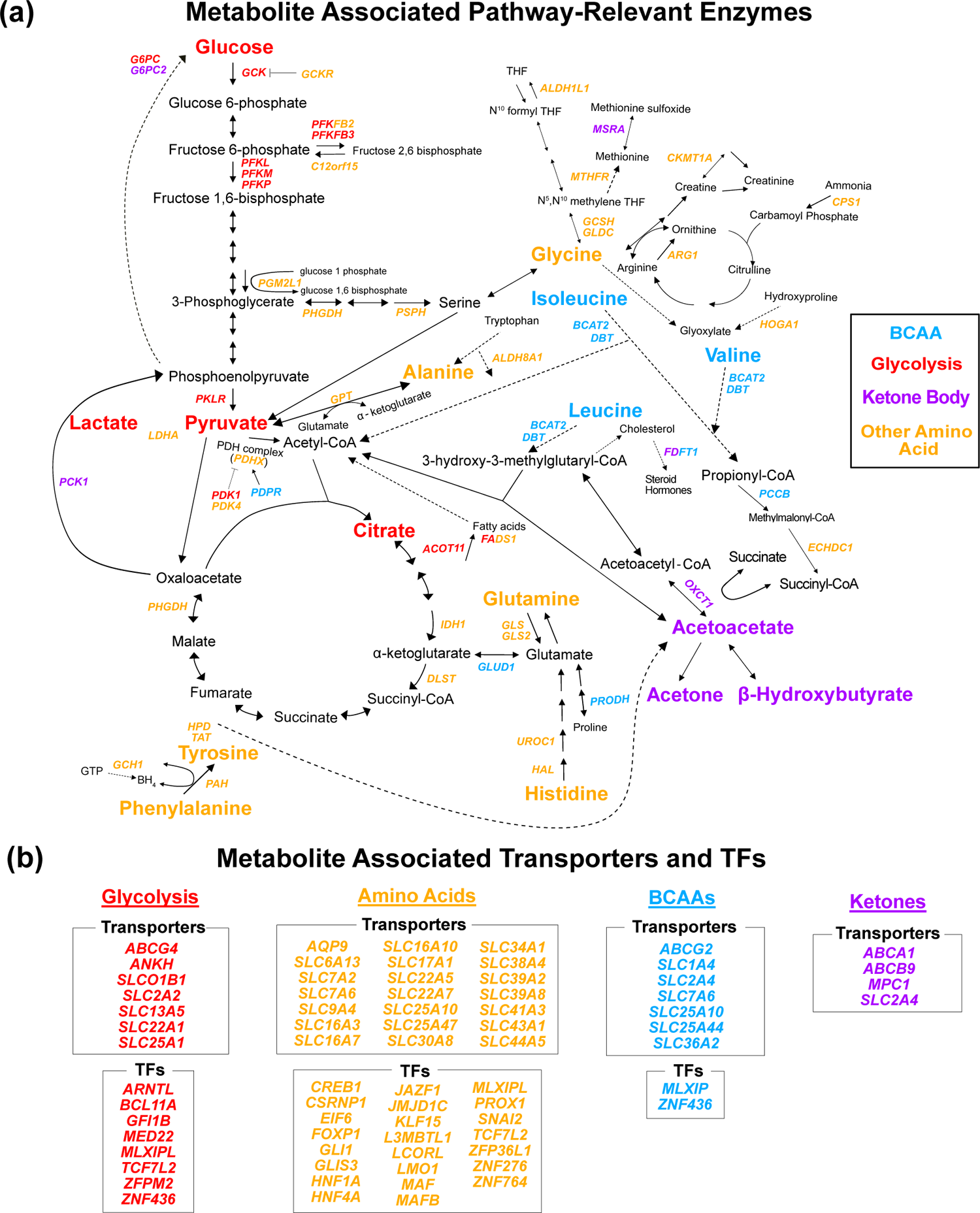
Gene annotation of metabolite GWAS hits. Each gene is colored based on the biochemical group with the most associated metabolites (P < 1e-4). If multiple biochemical groups are tied for the most associations for a given gene, they are all shown. **a.** Expanded pathway diagram with all genes (italicized) that encode pathway-relevant enzymes and were a metabolite GWAS hits. **b.** List of all genes of the metabolite GWAS hits that encode transporters and TFs. There were 69 metabolite GWAS hits that are not shown. Of these, 60 were annotated with genes assigned to the gene type general cell function (14 of these 60 were related to lipid function), and 9 were assigned to a gene of unknown function or that did not have any genes nearby (see Methods).

Next we sought to understand which genes and subpathways were most relevant to each biochemical group. We assigned each gene to the biochemical group with the most associated metabolites (Supplementary Table S4; see Methods). Genes were largely assigned to the group whose relevant biology was nearest the protein encoded by the gene. For example, *BCAT2* encodes an enzyme responsible for the first step in the breakdown of all three BCAAs and was assigned to the BCAA group. *OXCT1* encodes an enzyme responsible for the conversion of acetoacetyl-CoA to the ketone body acetoacetate and was assigned to the Ketone Body group. Similarly, *SLC7A9* encodes a protein that transports amino acids and was assigned to the Other Amino Acid group, while *TCF7L2* is a TF assigned to the Glycolysis group and involved in blood glucose homeostasis. These results confirm that these variants are affecting known trait-relevant biology and reflecting the local structure of these pathways.

Interestingly, a large fraction of the genes involved in trait-relevant biology were genome-wide significant hits for at least one of the 16 metabolites. Specifically, of the 139 total genes encoding enzymes in the pathway diagram for these metabolites (Supplementary Figure S4), 51 genes had at least one GWAS hit. In the followup ancestry-inclusive analysis, we identified 41 additional hits not found in the European-only GWAS, including associations at 7 additional pathway-relevant genes (Supplementary Table S5). This highlights the potential for large-scale, ancestry-inclusive GWAS to discover more biochemically-relevant associations among these traits. Together, these findings suggest that GWAS reflect, and have the potential to illuminate, the complex biochemical pathways interconverting these metabolites.

### Investigating the mechanisms of pleiotropy in trait pairs

Given the overlap between the biology of these metabolites and their hits, we next sought to understand the molecular causes of pleiotropy in trait pairs. We found 26 genetically correlated metabolite pairs at a local false sign rate < 0.005. For example, alanine and its strongest genetic correlation partner, isoleucine, share a genetic correlation of *r_g_* = 0.52 (P = 9e-23). Similarly, plotting the effects of the 213 GWAS variants on these two traits indicates a strong positive correlation (Figure 4a). Nonetheless, we noted several outlier loci, including rs370014171 (*PDPR*) and rs77010315 (*SLC36A2*), which have strong discordant effects. We were intrigued to understand why these two variants had discordant effects on alanine and isoleucine relative to their overall positive genetic correlation, while the majority of other variants had concordant effects.

Outlier variants are appealing case studies for understanding the molecular basis of pleiotropy because they affect traits in an exceptional way. Thus, we reasoned that understanding large-effect variants inconsistent with the global genetic correlation would reflect interesting biology relevant to the traits. For example, the proteins encoded by *PDPR* and *SLC36A2* are both located between alanine and isoleucine in the biochemical pathway (Figure 4b). This suggests that where variants act in the pathway may influence the direction of effect they have on metabolites. To better understand how these two variants affect alanine and isoleucine and explain their outlier behavior, we examined their effect size and direction in the context of their location in the pathway. We then used the variants’ metabolite associations to develop candidate mechanisms for how each variant could be jointly influencing the levels of these metabolites.

**Figure 4:**
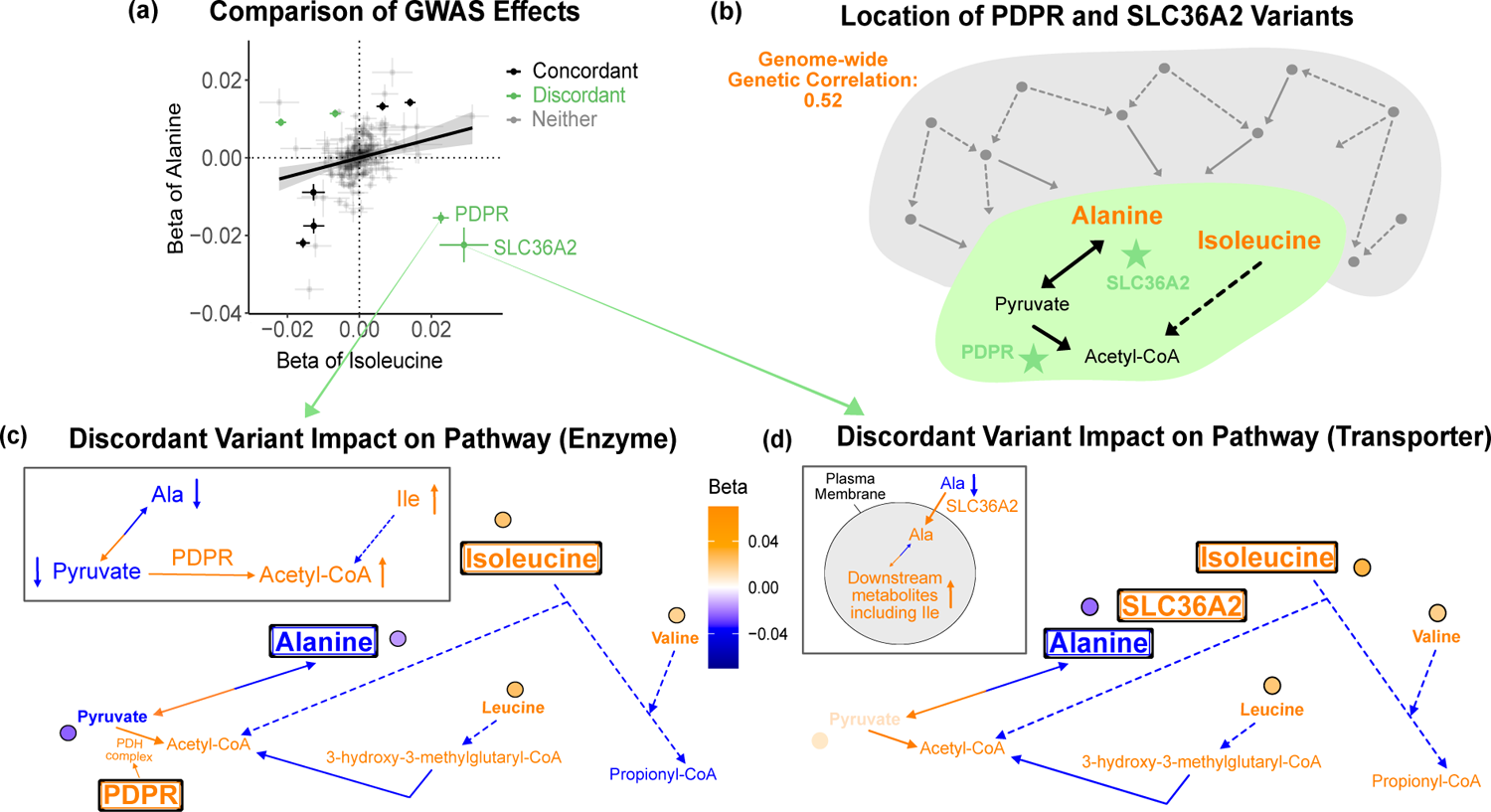
Discordant variant analysis. **a.** Comparison between the effects on alanine levels versus the effects on isoleucine levels for the 213 metabolite GWAS hits. This highlights two discordant variants: rs370014171 (PDPR) and rs77010315 (SLC36A2). Variants without an association P < 1e-4 in both metabolites are labeled “Neither”. **b.** Graphical representation of where these two discordant variants act in the pathway, represented by green stars, relative to other upstream variants driving the positive genetic correlation. Below are the hypothesized mechanisms explaining the GWAS results in relevant metabolites for each of these discordant variants. Data are shown in circles with the coloring corresponding to the effect (beta) of that variant on that metabolite. A black outline represents an association with P < 1e-4. Orange text and arrows represents a hypothesized increase (direction, not magnitude) in flux and blue corresponds to a decrease. **c.** Results for rs370014171 near the gene PDPR which encodes a protein that activates the conversion of pyruvate to acetyl-CoA. **d.** Results for rs77010315 in the gene SLC36A2 which encodes a small amino acid transporter.

As an illustration, we first consider variant rs370014171. This variant was assigned to gene *PDPR* because it was the second closest gene, the closest pathway-relevant enzyme, and within 100 kb (12.3 kb to its gene boundaries). PDPR activates the enzyme that catalyzes the conversion of pyruvate to acetyl-CoA (Figure 4c; Supplementary Figures S6 and S7). A candidate mechanism for this variant, supported by the effect size and direction for the 16 metabolites where relevant, is that it increases PDPR activity. This would lead to increased conversion of pyruvate to acetyl-CoA and thus decreased pyruvate (*β* = −0.023 SDs, P = 3e-20). To compensate for the subsequent decreased pyruvate levels, there would be increased conversion of alanine to pyruvate causing a decrease in alanine. In response to the increased acetyl-CoA, there would be decreased breakdown of metabolites normally catabolized for its production, including isoleucine, resulting in an increase in isoleucine levels. Thus, this variant has an opposite effect on alanine and isoleucine, despite their overall positive genetic correlation, likely because it affects the activity of an enzyme that acts in the pathway *between* the pair of metabolites. As expected due to the high correlation between the levels of the three BCAAs, this variant is also a discordant variant for alanine with valine (*r_g_* = 0.51, P = 2e-21), and alanine with leucine (*r_g_* = 0.49, P = 1e-16).

As a second example, variant rs77010315 is a missense variant in *SLC36A2*. *SLC36A2* encodes a transporter for small amino acids such as alanine (Figure 4d; Supplementary Figures S8 and S9). A candidate mechanism explaining the observed metabolite associations in our data and outlier behavior for this variant is that it increases transport of alanine into cells by SLC36A2. This would result in a decrease in levels of alanine in the blood, but an increase of alanine in cells. This additional intracellular alanine would then allow for increased conversion of alanine to pyruvate, thereby increasing levels of downstream metabolites in the blood, including isoleucine. Thus, this variant has an opposite effect on alanine and isoleucine, despite their overall positive genetic correlation, but in this case because it affects biology between the metabolites at the transporter level.

### Quantifying global properties of molecular pleiotropy

Based on these results, we hypothesized that the two variants described above, and others like them, exhibit outlier behavior because they affect biology *between* the two metabolites (Figure 5a). We consider biology “between” a given pair of metabolites as the shortest realistic biochemical path converting one to the other, and any alternative paths of reasonably similar distance and likelihood (see Methods for details). Genetic correlation reflects the direction of effect that most associated variants have on two traits. However, when two metabolites are biologically near each other, the region containing “between” biology is relatively small, such that only a minority of variants directly affect the “between” region.

**Figure 5:**
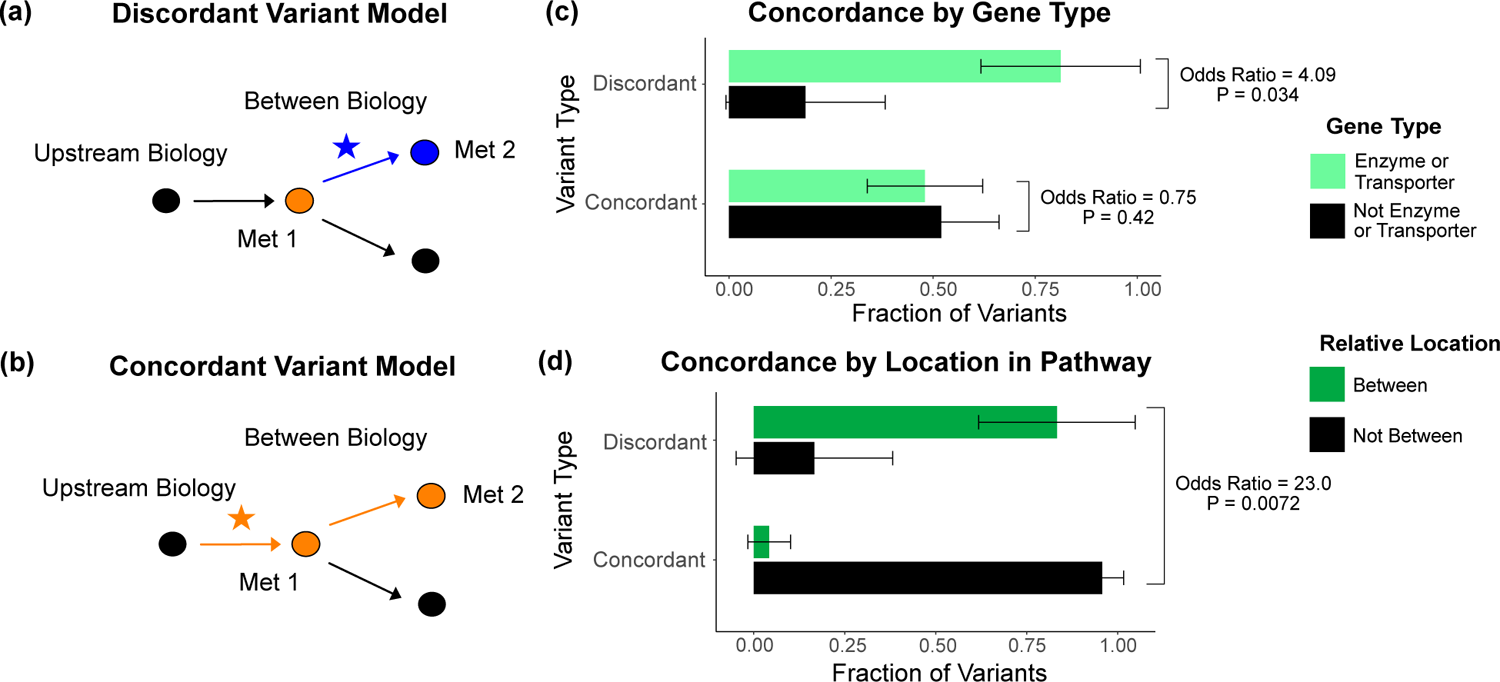
Characterization of discordant and concordant variants. **a.** Proposed model for the mechanism of a discordant variant. This example is for a discordant variant that has opposite effect directions on a pair of metabolites with a positive overall genetic correlation because it affects biology between them. **b.** Proposed model for the mechanism of a concordant variant. This example is for a concordant variant that has the same effect direction on a pair of metabolites with a positive overall genetic correlation because it affects biology upstream both metabolites. **c.** Fraction of the discordant and concordant variants that have a pathway-relevant enzyme or transporter gene type annotation versus those with a different gene type annotation. Discordant variants are enriched for the gene types of pathway-relevant enzyme or transporter, as would be expected in the model of discordant variants generally affecting biology between metabolites. **d.** Fraction of the discordant and concordant variants annotated with a pathway-relevant enzyme that affect biology between versus not between their significant metabolite pairs. Significance tests were performed using Fisher’s Exact method and the plotted SEs are from 95% CI calculated by binomial sampling variance.

Thus, we hypothesized that the genetic correlation of two biologically-related metabolites mostly reflects the effects of variants upstream or downstream of the metabolites, masking the effects of those between. We developed an analogous hypothesis that variants affecting biology upstream or downstream of the two metabolites have concordant effects (Figure 5b). While less common, the overall genetic correlation for two biologically-related metabolites can also be negative due to factors such as feedback loops. In this case, variants acting between the two metabolites would have the same direction of effect on both metabolites, making them discordant with the negative overall genetic correlation (Supplementary Figure S10).

To evaluate these models, we defined outliers based on the consistency of their effects with the overall LDSC genetic correlation. If a variant had an effect direction opposite the overall LDSC genetic correlation in at least one significant metabolite pair (P < 5e-8 in one, P < 1e-4 in the other), it was classified as “discordant”. For example, a discordant variant for a metabolite pair with a positive genetic correlation would have a negative association in one of the metabolites and a positive association in the other. If a variant had an effect direction consistent with the overall genetic correlation for its significant metabolite pairs, it was classified as “concordant”. Variants without multiple associations, or where associated traits were not significantly genetically correlated, were classified as “neither”. In total, of the 62 metabolite GWAS hits that had at least one significant metabolite pair, we found 26 total discordant variant-metabolite pairs across 14 variants (Supplementary Table S6).

We then investigated overall properties of discordant variants relative to concordant ones. We discovered that discordant variants are more likely to affect genes encoding enzymes and transporters than all other genes types, including TFs, general cell function genes, and those of unknown function (Odds Ratio = 4.09, P = 0.034; Figure 5c). This is in contrast to concordant variants, which do not show an enrichment for enzymes and transporters relative to other gene types (Odds Ratio = 0.75, P = 0.42). These observations are consistent with our model that discordant variants tend to affect biology between relevant pairs of metabolites since TFs and general cell function genes generally act outside these metabolic pathways. Thus, they are more likely to affect biology upstream or downstream of both metabolites. In addition, for variants affecting pathway-relevant enzymes, where the location in the pathway that the variant is acting relative to the metabolites is clear, we were able to directly test our hypothesis. We found that discordant variants affecting pathway-relevant enzymes are much more likely to act between, rather than upstream or downstream, the metabolites for which they are discordant (Odds Ratio = 23.0, P = 0.0072; Figure 5d).

We then sought to extend this finding by developing a model contrasting the effects of all variants affecting between versus outside biology at a pathway and genome-wide level (Figure 6a). In aggregate, this model predicts that pathways overlapping biology between two metabolites will have a local genetic correlation opposite that of nearby adjacent pathways and that the magnitude of both will exceed that of the global polygenic background. As a case study, we focused on alanine and glutamine, which have a weak positive overall genetic correlation (*r_g_* = 0.16, P = 0.08; Figure 6b; Supplementary Figure S11). We then ran BOLT-REML [29] on variants within 100 kb of genes in each pathway and estimated the corresponding local genetic correlations (see Methods).

**Figure 6:**
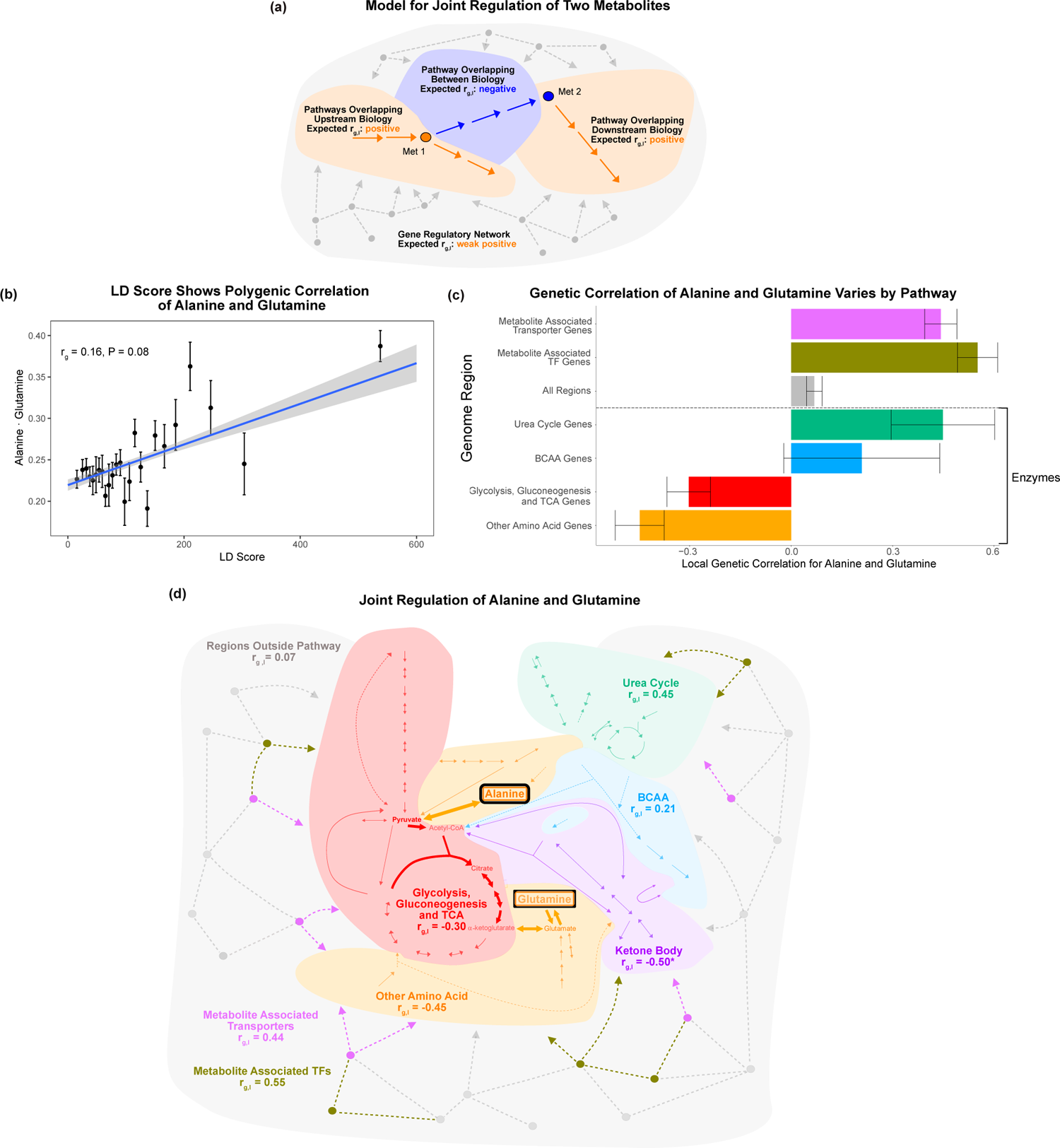
Local genetic correlation. **a.** Model of expected local genetic correlation direction, with contrasting effects of variants affecting “between” versus outside biology at a pathway and genome-wide level. **b.** LD Score showing the polygenic correlation of alanine and glutamine. For the x axis, LD Scores were binned into 25 bins. The y axis shows the mean and SE within each bin. **c.** Results for the local genetic correlation of alanine and glutamine for variants within 100 kb of genes in each pathway. Standard errors are shown. Genesets listed below the dotted line include only enzymes and are considered pathway-relevant enzymes for these metabolites. Summary statistics for BOLT-REML and other methods can be found in Supplementary Table S7. **d.** Pathway diagram showing the pathways included in the local genetic correlation analysis and the positioning of their genes relative to alanine and glutamine. *Ketone Body Genes were omitted from panel c because the limited number of genes meant they failed to robustly converge. All arrows and nodes in the gray section are hypothetical and shown for illustration purposes.

We found that the local genetic correlations around genes in the Glycolysis, Gluconeogenesis and Citric Acid Cycle Pathway and around genes in the Other Amino Acid Pathway were negative (Figure 6c). Both of these pathways encompass genes affecting biology between alanine and glutamine (Figure 6d). In striking contrast, nearby pathways, such as the Urea Cycle, had a positive local genetic correlation for these metabolites (*r_g,l_* = 0.45; P = 0.003). Similarly, we found that regions overlapping genes encoding metabolite associated transporters and TFs had strong positive genetic correlations consistent with their shared role in the upstream regulation of these two traits (*r_g,l_* = 0.44, P = 1e-20; *r_g,l_* = 0.55, P = 2e-20). All genes outside the core pathways had a weak positive genetic correlation, perhaps reflecting that they are embedded in the global gene regulatory network (*r_g,l_* = 0.068; P = 0.003). Our findings were broadly consistent using individual level data with Haseman-Elston regression [30], and summary statistics with *ρ*-HESS [3], stratified LD score regression [31] and a non-parametric Fligner-Killeen variance test (see Methods; Supplementary Table S7). These results support the model that variants affecting biology between the metabolites frequently contrast with the contributions of upstream and downstream pathways. This emphasizes that the heterogeneity in genetic effects reflecting local biology shared by the traits can be masked in the global genetic correlation. In addition, these results offer biological intuition for interpreting genetic correlation of molecular traits at a pathway and genome-wide level.

### Using metabolites to understand the mechanism of a disease-associated variant

Motivated by the interpretability of these results, we applied this logic to understand the mechanism underlying disease-associated variants. We developed an example model for a disease-associated variant impacting a relevant subpathway and consequent metabolite levels in a way that is consistent with disease etiology (Figure 7a). To apply this model to our data, we considered variants that were annotated with pathway-relevant enzymes and associated with increased risk for coronary artery disease (CAD) [32, 33]. The strongest variant we identified, rs61791721, was assigned the nearest pathway-relevant enzyme gene, *PCCB*. *PCCB* encodes a protein that catalyzes the conversion of propionyl-CoA to succinyl-CoA at the intersection of BCAA and fatty acid oxidation (Supplementary Figure S12).

**Figure 7:**
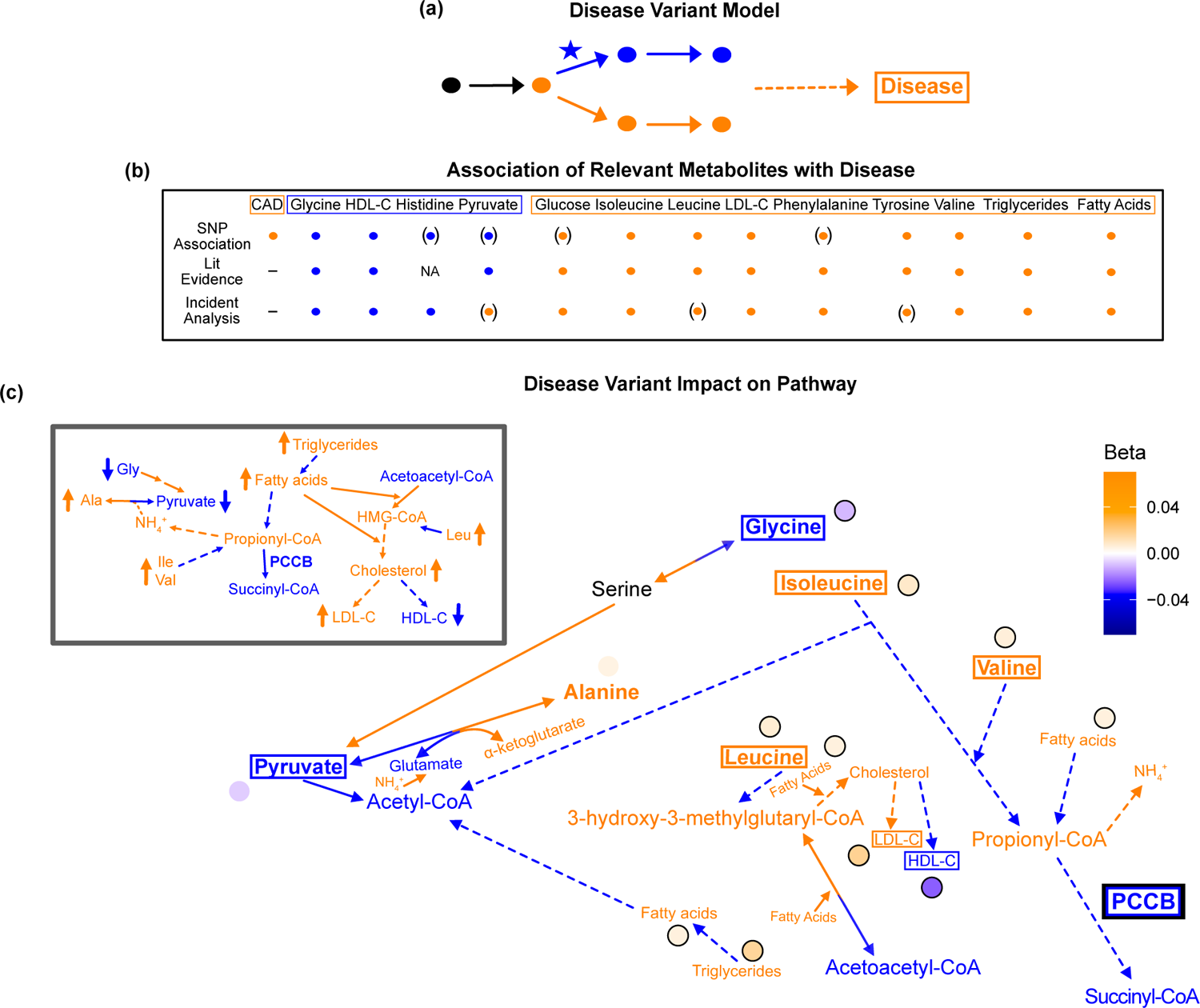
Pathway impact and pathology of example disease GWAS hit. **a.** Proposed model for the impact of a disease hit on a relevant pathway, contributing to an increased risk in the disease. **b.** This variant is associated with an increase in levels of metabolites that have been implicated with increased risk of CAD, and a decrease in the levels of metabolites that have been implicated with decreased risk. Parentheses indicate nonsignificant associations, “NA” indicates no evidence was found, and “-” indicates a placeholder because CAD is being compared with itself. **c.** Results for rs61791721 with gene assignment PCCB which encodes a protein that catalyzes the conversion of propionyl-CoA to succinyl-CoA. The hypothesized mechanism is that the variant is decreasing the activity of PCCB, resulting in the above metabolite associations. Ammonium is represented by its chemical formula (NH^+^).

We combined results from the literature and incident analysis to understand the association of relevant metabolites with CAD (Supplementary Figure S13). We then compared these to the effects of this variant on these metabolites (Figure 7b). In this analysis we included high density lipoprotein cholesterol (HDL-C), low density lipoprotein cholesterol (LDL-C), total fatty acids, and total triglycerides, due to the extensive evidence implicating their association with CAD, and because they are directly adjacent to the biology of the other 16 metabolites. Consistent with the metabolites’ corresponding risk for CAD, this *PCCB* variant was negatively associated with glycine and HDL-C, and positively associated with isoleucine, leucine, valine, tyrosine, total fatty acids, total triglycerides and LDL-C (P < 1e-5; Supplementary Tables S8 and S9).

This PCCB variant has been associated with CAD in multiple prior GWAS [32, 33], yet neither the gene this variant affects nor the mechanism explaining this association are known. However, this variant affects many metabolites associated with CAD in a direction consistent with increased risk. Thus, we can begin to understand why this variant is associated with CAD by understanding the pleiotropic effects of this variant on the metabolites.

The hypothesized mechanism resulting in this pathogenic constellation of metabolite effects is that the variant decreases PCCB activity, resulting in lower levels of succinyl-CoA and increased propionyl-CoA (Figure 7c; Supplementary Figure S14). The increased propionyl-CoA would result in excess ammonium being produced, and because alanine is a reservoir for nitrogen waste, this would increase conversion of pyruvate to alanine to capture the toxic ammonium [34, 35]. More glycine would be broken down in response to the decrease in pyruvate levels, decreasing glycine levels. Conversely, the increased levels of propionyl-CoA mean less valine, isoleucine, fatty acids, and thus triglycerides, would need to be broken down, resulting in an increase in their levels. This increase in fatty acids may stimulate the activity of 3-hydroxy-3-methylglutaryl-CoA (HMG-CoA) reductase and synthase, resulting in an increase in HMG-CoA and cholesterol [36, 37]. Increased HMG-CoA would lead to increased leucine because less leucine would need to be broken down to produce HMG-CoA, while increased cholesterol would lead to an increase in LDL-C and a decrease in HDL-C. Therefore, this variant is potentially associated with CAD because it is decreasing PCCB activity, resulting in myriad deleterious downstream metabolic consequences.

This example demonstrates that we can begin to dissect the molecular basis underlying disease GWAS hits by understanding the mechanism of relevant pleiotropic effects on metabolites. In addition, the pathways implicated by this analysis can also be independently prioritized as potentially playing an important role in cardiometabolic disease by leveraging the molecular basis of genetic correlation discussed in Figure 6. For example, alanine and glutamine have opposite associations with CAD and type 2 diabetes despite having an overall positive phenotypic correlation [38, 39]. This suggests that the pathways described above with a negative local genetic correlation for alanine and glutamine are likely relevant to the molecular basis of these diseases. Thus, understanding the molecular basis of pleiotropy and genetic correlation of metabolites can improve our understanding of the variants and pathways contributing to complex disease biology.

## Discussion

In this work, we investigate the joint effects of pleiotropic variants on 16 biologically-related metabolites in the context of their biochemical pathways. We build on prior studies examining the genetic architecture of metabolites by characterizing the genes and mechanisms through which variants affect these metabolites, and find a strong enrichment for genes encoding pathway-relevant enzymes and transporters. Our results offer biological intuition for the interpreting genetic correlation of molecular traits at a pathway and genome-wide level.

We demonstrate the effects of variants acting on biology between metabolites often contrast substantially with the contributions of upstream and downstream pathways, as well as the polygenic background. Perhaps paradoxically, while the overall genetic correlation between two traits provides a global view of shared effects, the genes that are directly involved in the traits’ core biology are most likely to have divergent effects. We show that one explanation of this is the substantial outlier contributions from variants acting directly between metabolites of interest. We anticipate that further mechanisms, such as context-specific variant effects and differential regulation by peripheral genes, will be discovered in future studies.

In addition, we show specific examples of candidate molecular mechanisms explaining the association of variants with multiple biologically-related metabolites. These include associations at *PDPR*, *SLC36A2*, and *PCCB*, where we show that the direction and magnitude of their effects is consistent with metabolite biochemistry and disease etiology. These proposed molecular mechanisms enable biological prioritization of interesting candidates for future post-GWAS *in vitro* studies. Overall, these results suggest specific genetic and molecular underpinnings of complex disease variants, and provide a roadmap for further discovery through the interpretation of pleiotropic variant effects on disease-relevant metabolites.

In this work, we focus on metabolites clustered at the intersection of amino acid catabolism, glycolysis, and ketone body metabolism. However, the approaches and results from this paper have the potential to reveal novel insights into genetic effects on many biochemical pathways and molecular traits. In addition, integrating this work with proteomic and intermediate metabolomic data will offer additional evidence to develop and support these hypothesized mechanisms. These data may also clarify the relevance of additional mechanisms – such as buffering, feedback, and kinetics – in controlling the plasma levels of these metabolites. Finally, expanding the sample size and diversity of ancestries included in future GWAS, measurements of which are currently underway on the Nightingale platform and others, will increase power to detect novel findings such as important associations for variants with low allele frequency.

One limitation of this study is that the metabolites were measured in the blood, while most of the relevant biology and pathways occurs within cells in various tissues throughout the body. Thus, we anticipate extensions of this work to include biomarker measurements from additional cell types and tissues, such as urine, saliva, biopsy samples, and *in vitro*-differentiated cells. Further, longitudinal analysis of relevant disease cohorts will allow insights into disease progression and subtyping.

In conclusion, this work underscores the potential of unifying biochemistry with genetic data to understand the molecular basis of complex traits and diseases and the mechanism through which variants impact these traits.

### Online Methods

#### Population Definition

We defined our GWAS population as a subset of the UK Biobank [40]. For our cohort, we use the individuals for which Nightingale plasma metabolite data was available after filtering based on trait QC characteristics (see “Trait QC and Covariate Adjustment”). We then filtered individuals by the following QC metrics:

1. Not marked as outliers for heterozygosity and missing rates (`het_missing_outliers` column)
2. Do not show putative sex chromosome aneuploidy (`putative_sex_chromosome_aneuploidy` column)
3. Have at most 10 putative third-degree relatives (`excess_relatives` column).
4. No closer than second degree relatives.

From these, we defined 3 cohorts: White British, non-British White, and everyone. We identified White British individuals using the `in_white_British_ancestry_subset` column in the sample QC file. We identified non-British White individuals through self-identification as White, excluding individuals marked as `in_white_British_ancestry_subset` (n = 30,116 who passed QC metrics 1-4 above). As was done for the White British in the initial UK Biobank study design [40], we identified global principal components of the genotype data, and then defined ancestry clusters using `aberrant` with the strictness parameter *λ* = 20. Non-British White individuals who were outliers for any of projected principal component pairs PC1/PC2, PC3/PC4, and PC5/PC6 were excluded (n = 25,137 remaining). We performed our first GWAS on the set of individuals in these White British and non-British White cohorts.

The combination of the two sources of European and White British ancestry individuals resulted in a total of 433,390 European ancestry individuals in UK Biobank, of whom 94,464 had available quality controlled Nightingale data. Our main goal for this study was to understand general principles of genetic architecture, which are not expected to vary among human populations, and thus in the main analysis we excluded non-European individuals on the basis of power and concerns about structure confounding. However, this analysis is significantly limited by the allele frequency differences between populations, and we sought to develop an alternative, inclusive strategy that did not rely on self-identity.

For the ancestry-inclusive analysis, we performed the same QC steps 1-4 without filtering individuals on the basis of self-identified race/ethnicity or on the basis of their ancestry PC outlier status. This resulted in a total of 98,189 individuals for the GWAS. This was inspired by recent “mega-analysis” studies [41].

### Metabolomics Data Generation

The metabolomics data was generated by Nightingale Health using a high-throughput NMR-based platform developed by Nightingale Health Ltd. Randomly selected EDTA non-fasting (average 4h since last meal) plasma samples (aliquot 3) from approximately 120,000 UK Biobank participants were measured in molar concentration units. The included participants are therefore meant to be representative of the 502,543 participants in the full cohort. The measurements took place between June 2019 and April 2020 using six spectrometers at Nightingale Health, based in Finland. The Nightingale NMR biomarker profile contains 249 metabolic measures from each plasma sample in a single experimental assay, including 168 measures in absolute levels and 81 ratio measures. The biomarker coverage is based on feasibility for accurate quantification in a high-throughput manner and therefore mostly reflects molecules with high concentration in circulation, rather than selected based on prior biological knowledge. Additional details about the data generation can be found at https://biobank.ndph.ox.ac.uk/showcase/ukb/docs/nmrm_companion_doc.pdf.

### Trait Selection and Grouping

Sixteen metabolites were chosen from the available Nightingale metabolites based on their biochemical proximity, relevance to health and disease, and because the genes and enzymes involved in their metabolism are well-characterized. Specifically, we first filtered to the 168 metabolites that were not metabolite ratio measurements (n=81) because we wanted to focus on absolute metabolites levels. We then filtered out the lipids and lipoprotein measures, including cholesterol and fatty acids, because the complexity of their biochemistry make it difficult to map out the chemical reactions directly interconverting one to another, and because many of these metabolites have already been extensively studied in large GWAS [42, 20]. However, many of these are important metabolites in the discussion of cardiometabolic disease so we additionally ran GWAS for total triglycerides, total fatty acids, HDL cholesterol, and LDL cholesterol as part of the interpretation of the *PCCB* variant using the same pipeline as for the sixteen metabolites below.

Finally we removed remaining derived measures (such as total combined concentration of BCAA) and those primarily reflecting physiological conditions such as fluid balance (creatinine and albumin) and GlycA (inflammation). One exception to this filtering was the three ketone bodies (3-hydroxybutyrate, Acetone and Acetoacetate) which were included due to their proximity and clear direct interconversions connecting them to the metabolic pathways of the remaining amino acid and glycolysis-related metabolites. Metabolites were classified into four biochemical groups based on biochemical similarity. The three branched chain amino acids: valine, leucine and isoleucine were classified as “BCAA”, the remaining amino acids in the dataset: glycine, glutamine, tyrosine, phenylalanine and histidine were classified as “Other Amino Acid”, the three ketone bodies: 3-hydroxybutyrate, acetone and acetoacetate were classified as “Ketone Body” and the four metabolites in or immediately adjacent to glycolysis: glucose, pyruvate, lactate and citrate were classified as “Glycolysis”.

### Trait QC and Covariate Adjustment

Trait measurements were filtered to only include baseline samples then log-transformed. Outlier removal was performed by dropping any sample that had a metabolite level greater than 20 fold the interquartile range or greater than 10 fold below the median across all samples for that metabolite. PCA was run for the remaining samples and outliers were dropped using aberrant (lambda = 20) on the top two PCs [43]. Remaining log transformed measurements were adjusted for spectrometer, week, and weekday. Samples were subset to the GWAS population defined above resulting in 94,464 individuals for the European ancestry GWAS and 98,189 individuals for the ancestry-inclusive GWAS.

### GWAS

We performed GWAS in BOLT-LMM v2.3.2 [29] adjusting for sex, array, age, and genotype principal components 1-10 using the following command (data loading arguments removed for brevity):

`bolt --phenoCol= [Metabolite] \`

`--covarCol=sex \`

`--covarCol=Array \`

`--qCovarCol=age \`

`--qCovarCol=PC{1:10} \`

`--lmmForceNonInf \`

`--numThreads=24 \`

`--bgenMinMAF=1e-3 \`

`--bgenMinINFO=0.3`

The resulting GWAS summary statistics were then filtered to minor allele frequency greater than 0.01 and INFO score greater than 0.7 for further analyses (referred to as the Filtered Metabolite Sumstats). The LDSC munge_sumstats.py script was then use to munge the data (referred to as the Munged Metabolite Sumstats) [2].

### GWAS Hit Processing

To evaluate GWAS hits, we took the Filtered Metabolite Sumstats and ran the following command using plink version 1.9 [44]:

`plink --bfile [] --clump [GWAS input file] --clump-p1 1e-4 --clump-p2 1e-4`

`--clump-r2 0.01 --clump-kb 1000 --clump-field P_BOLT_LMM --clump-snp-field SNP`

We greedily merged GWAS hits across the 16 metabolites located within 0.1 cM of each other and took the SNP with the minimum p-value across all merged lead SNPs. In this way, we avoided potential overlapping variants that were driven by the same, extremely large, gene effects. This resulted in 213 lead GWAS variants, referred to as the metabolite GWAS hits.

For the ancestry-inclusive analysis, we used the European ancestry LD matrix as European-ancestry individuals were the overwhelming majority in the study. Here, we identified 238 lead GWAS variants.

### Gene and Gene Type Annotation

We defined all genes in any GO [45, 46], KEGG [47], or Reactome MSigDB [48, 49] pathway as our full list of putative genes (in order to avoid pseudogenes and genes of unknown function). We initially extended genes by 100 kb (truncating at the chromosome ends) and used the corresponding regions, overlapped with SNP positions, to define SNPs within range of a given gene. Gene positions were defined based on Ensembl 87 gene annotations on the GRCh37 genome build. We then performed manual curation using GeneCards [50] to validate gene assignments and prioritize a single gene per SNP. Gene boundaries for genes encoding pathway-relevant enzymes in KEGG were extended up to 500 kb and assigned to a variant if the gene was biologically relevant to the metabolites the variant was significant in. If there were multiple genes within 100 kb of the variant, then gene assignments were made based on the following priority order: any genes encoding a pathway-relevant enzyme, genes encoding transporters, genes involved in translation/transcription regulation (referred to as TF for brevity), any genes whose function is known. If there were multiple genes of the same gene type, then the assignment was made based on the relevance of the gene to the metabolites the variant was significant in, proximity of the gene to the variant, and, if applicable, any additional evidence in the literature (Oxford BIG [51] and Open Target Genetics [52, 53]). However, even for these cases where there was not high confidence in the exact gene assignment, for instance because there were multiple genes from the same gene family nearby, the top gene candidates all had the same gene type. Thus, because the major downstream analyses were designed in a way that only the gene type assigned to each variant mattered, the accuracy of the exact gene assignment should not affect the findings. If no genes with known function were within 100 kb of the variant then the window was extended up to 200 kb. The distance of a variant to a given gene was defined as the number of base pairs from the variant to the closer of the start or end of the gene boundaries or was set to 0 if the variant was within the gene boundaries.

We classified each gene using GeneCards [50] into one of five gene types: pathway-relevant enzyme, transporter, TF, general cell function and unknown. Genes encoding enzymes that catalyze a reaction in or adjacent to the direct synthesis or degradation of one of the 16 metabolites were defined as pathway-relevant enzymes using manual curation from GO, KEGG, REACTOME and Stanford’s Human Metabolism Map [54], in addition to GeneCards. Genes encoding known transporters were classified as transporters. Genes involved in translation/transcription regulation were classified as TF. Genes whose function is known but not already classified as a pathway-relevant enzyme, transporter, or TF, were classified as “general cell function”. Genes with unknown function or if there were no genes within 200 kb of the SNP were classified as “unknown”. See S3 for each metabolite GWAS hit’s gene and gene type annotations.

Gene type enrichments were calculated with a Poisson rate test. The baseline was the total of the GWAS hits among the 1.95 Gb of the genome within 100 kb of a gene in any pathway, and the test was performed with the number of GWAS hits within 100 kb of each pathway of interebst. There were 2.8 GWAS hits per megabase within 100 kb of a pathway-relevant enzyme versus 0.1 GWAS hits per megabase among all genes (25-fold enrichment, Poisson rate test P < 2e-16). There were 0.58 GWAS hits per megabase within 100 kb of a transporter versus 0.1 GWAS hits per megabase for all genes (5.2-fold enrichment, Poisson rate test P = 9e-16). We also repeated this analysis using closest genes rather than assigned genes, which allowed us to use a Fisher exact test (as each variant has a single closest gene). This resulted in a 27-fold (P < 2e-16) enrichment for pathway-relevant enzymes and an 11-fold (P < 2e-15) enrichment for transporters, respectively.

For TF enrichments, we used TF-Marker [55] to annotate tissue-specific marker gene TFs. We considered “TF” (n = 1316) and “TFMarker” (n = 18) genes as relevant genes, and TF Pmarker (n = 1424) genes as putatively relevant. We considered enrichment among the 628 genes not associated with cancer or stem cell biology (of which 267 are putative) as our set of tissue-specific TFs for downstream analysis. We consider our background in all cases to be our total GWAS hits number (n = 213) compared to the effective genome size (2.86 Gb). In specifically this TF set, we observed a 5.96-fold enrichment over the genome wide background (0.44 GWAS hits per megabase, p = 4e-6) among relevant gene bodies and 2.50-fold enrichment (0.19 hits/Mb, P = 0.0007) within 100 kb of a relevant gene, which was comparable for putatively relevant genes (4.85-fold and 2.86-fold, respectively). This was substantially higher than that of all genes in the genome (1.69-fold within gene bodies and 1.36-fold within 100kb of genes in any pathway) and comparable to that of all TFs regardless of their function in cancer or stem cells (4.82-fold within gene bodies and 2.18-fold within 100kb).

We next filtered tissue-specific TFs to those acting in Liver (28 relevant and 30 putative), Kidney (25 relevant and 20 putative), or Pancreas (14 relevant and 3 putative). Kidney and Pancreas TFs had no more than 1 GWAS hit each and were excluded for these analyses. For Liver TFs, we observed a 18-fold enrichment (1.3 GWAS hits per megabase, P = 0.0057) within gene bodies and a 7.6-fold enrichment (0.56 hits/Mb, P = 0.002) within 100kb of genes. Results were similar when removing the cancer and stem cell filter (1.23 hits/Mb and 0.51 hits/Mb respectively) and dropped slightly when further including putatively relevant TFs (0.71 hits/Mb and 0.48 hits/Mb). Together, this suggests that liver marker TFs are specifically enriched for variants affecting our metabolite levels.

### HESS Trait Heritability and Pathway Enrichments

We ran HESS [56] using the following commands:

`hess.py --local-hsqg {\filtsumstats} --chrom {chrom} \`

`--bfile 1kg_eur_1pct_chr{chrom} --partition EUR/fourier_ls-chr{chrom}.bed \`

`--out {Metabolite}_step1`

`hess.py --prefix {Metabolite}_step1 --out {Metabolite}_step2`

Where `1kg_eur_1pct_chr{chrom}` were downloaded from: https://ucla.box.com/shared/static/l8cjbl5jsnghhicn0gdej026x017aj9u.gz and `EUR/fourier_ls-chr{chrom}.bed` were downloaded from:

https://bitbucket.org/nygcresearch/ldetect-data/src/ac125e47bf7f/?at=master

We intersected the resulting heritability estimates per LD block with gene lists from each pathway (see Local *ρ*-HESS; within 100 kb of the gene boundary was used as the tested window) and calculated the total heritability within the pathway as the sum of the heritabilities across LD blocks and the variance of the heritability within the pathway as the sum of the variances within each LD block. Overall, this gave a per-pathway estimate. We generated genome-wide estimates of heritability as well as heritability estimates for the subset of the genome nearby any coding gene in MSigDB as background controls from which to estimate the heritability enrichments, and used the coding gene numbers for reporting as they are more conservative.

### LDSC Genetic Correlation

LD Score regression [2] was used to generate genetic correlation estimates. The following command was used:

`ldsc.py --rg {\mungsumstats} --ref-ld-chr eur_ref_ld_chr`

`--w-ld-chr eur_w_ld_chr`

`eur_*_ld_chr` were downloaded from https://data.broadinstitute.org/alkesgroup/LDSCORE/.

### Mendelian Randomization

The Rücker model selection framework was applied. Briefly, MR was run with inverse-variance weighted (IVW) and MR-Egger with fixed and random effects, and selection between different methods for results to present was based on the goodness-of-fit and heterogeneity parameters for the individual MR regressions as previously described [57, 58].

### Discordant Variant Analysis

All pairwise combinations of LDSC Genetic Correlation (as described above) were performed for the 16 metabolites. Pairs were filtered to those that had a genetic correlation significantly different than 0 using ashR [59] with a local false sign rate of 0.005. We then annotated all metabolite GWAS hits with pairs of metabolites for which the variant had a P < 1e-4 association with both metabolites and a P < 5e-8 association with at least one, defined as significant metabolite pairs. A variant was classified as “discordant” if it had the same effect direction in both metabolites of at least one significant metabolite pair that had a negative global genetic correlation, or if it had opposite effect directions in the two metabolites of at least one significant metabolite pair that had a positive global genetic correlation. 14 variants of the 62 that had at least one significant metabolite pair were classified as discordant. Variants that had the same set of effect directions as the sign of the global LDSC genetic correlation for all of its significant metabolite pairs were classified as “concordant”. Variants that had no significant metabolite pairs were classified as “neither”.

The “between” region for a given pair of metabolites was defined as the shortest realistic biochemical path converting one to the other, and any alternative paths of reasonably similar distance and likelihood. Genes were defined as acting between a given metabolite pair either if they encoded an enzyme catalyzing a reaction in the “between” region defined above or if they encoded a transporter that primarily transports either of the two metabolites themselves or an intermediate metabolite in the “between” region. Variants were defined as acting between a given metabolite pair if the gene they affect was defined as between. Pathways were defined as between a given metabolite pair if many of the genes defined as between the metabolites were part of the pathway or if many of the genes in the pathway were defined as between. Note that even if a pathway is defined as “between”, not all genes in the pathway will always be between and vice versa; however, this is likely to only make the resulting differences in genetic correlation for “between” vs not “between” pathways more conservative.

### Local ***ρ***-HESS

We ran HESS [3] using the following commands:

`hess.py --local-rhog {Met1_sumstats}`

`{Met2_sumstats} --chrom {chrom} --bfile 1kg_eur_1pct_chr{chrom} \`

`--partition EUR/fourier_ls-chr{chrom}.bed --out {Met1_Met2}_step1 \ hess.py --prefix {Met1_Met2}_step1_trait1 \`

`--out {Met1_Met2}_step2_trait1 hess.py --prefix {Met1_Met2}_step1_trait2 \`

`--out {Met1_Met2}_step2_trait2 hess.py --prefix {Met1_Met2}_step1 \`

`--local-hsqg-est {Met1_Met2}_step2_trait1 {Met1_Met2}_step2_trait2 \`

`--num-shared 94464 \`

`--pheno-cor {gcov_int from LDSC genetic correlation for Met1_Met2} \`

`--out {Met1_Met2}_step3`

Where `1kg_eur_1pct_chr{chrom}` were downloaded from: https://ucla.box.com/shared/static/l8cjbl5jsnghhicn0gdej026x017aj9u.gz and `EUR/fourier_ls-chr{chrom}.bed` were downloaded from:

https://bitbucket.org/nygcresearch/ldetect-data/src/ac125e47bf7f/?at=master

We then used the local rho HESS results and estimated the local genetic covariance and correlation across all LD blocks overlapping pathway regions.

We defined the pathway regions based on gene boundaries of relevant genes in Supplementary Figure S4 as follows: “Other Amino Acid Genes” includes all genes colored orange, “Ketone Body Genes” includes all genes colored purple, “Glycolysis, Gluconeogenesis and TCA Genes” includes all genes colored red, and “Urea Cycle Genes” includes all genes colored green. “BCAA Genes” included all genes in KEGG_VALINE_LEUCINE_AND_ISOLEUCINE_DEGRADATION except *OXCT2*, *HMGCL*, *HMGCS1*, *HMGCS2*, *ACAT1*, *ACAT2*, *OXCT1*, *DLD*, *AGXT2*, *ABAT*, and *AACS* and also included *ECHDC1*. “All Regions Outside Pathway Genes” was defined as all LD blocks not overlapping any of the regions defined above. “Metabolite Associated TF Genes” and “Metabolite Associated Transporters Genes” were defined as all LD blocks overlapping any of TFs or Transporters respectively annotating the Metabolite GWAS Hits.

### Fligner-Killeen Variance Test

Rather than aggregating variant effects and estimating total genetic covariance and heritability per pathway, which is not robust to outlier effects, we additionally tried a non-parametric approach. Individual *r_g_* and *h*^2^ estimates for LD blocks were compared between the baseline (all coding genes) and the pathway of interest by listing all per-block genetic covariance scores and computing a Fligner-Killeen Variance Test within each pathway in R. This enables direct evaluation of genetic covariances between the pathways, at the cost of simultaneously capturing the enrichment of heritability and genetic covariance therein.

### BOLT-REML

Genotyped variants within 100 kb of genes in each pathway were aggregated, and the resulting matrices were tested using the following command in BOLT-LMM:

`bolt`

`--remove {non-European ancestry individuals}`

`--phenoFile={Technical-adjusted metabolites} \`

`--phenoCol=Ala \`

`--phenoCol=Gln \`

`--covarCol=sex --covarCol=Array --qCovarCol=age --qCovarCol=PC{1:10} \`

`--geneticMapFile=genetic_map_hg19_withX.txt.gz ‘# downloaded with bolt’ \`

`--numThreads=24 --verboseStats \`

`--modelSnps {pathway SNPs} \`

`--reml \`

`--noMapCheck`

Standard errors were as reported by BOLT-REML.

### Haseman-Elston Regression

Genotyped variants were pruned to MAF > 1% and approximate linkage equilibrium among individuals included in the GWAS using `--indep 50 5 0.5`. The resulting variants were used to construct genetic relatedness matrices (GRMs) that included the genotyped SNPs within 100 kb of genes in each pathway, and the resulting matrices were tested using the following command in GCTA:

`{GCTA} --HEreg-bivar {trait1} {trait2} --thread-num 16 --grm {GRM}`

Results using multiple GRMs (`--mgrm`) to jointly test all pathways were qualitatively similar outside of the genome-wide GRM, which no longer captured the within-pathway component.

### Stratified LD Score Regression

Analyses were performed as described in LDSC Genetic Correlation, except that rather than `eur_ref_ld_chr` as the reference LD Scores, instead LD Scores computed on variants within 100 kb of genes in each pathway were utilized.

### Disease Variant Analysis

The metabolite GWAS hits annotated with pathway-relevant enzymes were overlapped with significant hits for CAD, identifying the variant rs61791721 as the most significant variant [32, 33]. Incident coronary artery disease cases were defined among UK Biobank participants as those individuals who received a first diagnosis of myocardial infarction (MI) using the analytical MI model (field 42000) after the date of baseline assessment. Prevalent cases (individuals with a first diagnosis before date of assessment) were excluded. A Cox proportional hazard model was run with the technical-covariate-adjusted, log-transformed metabolite levels predicting incident MI status, adjusted for age, age^2^, age * sex, age^2^ * sex, and statin usage (defined based on a list of individual drug codes as previously described [7]). Effect sizes presented are based on the estimates from these models run independently for each metabolite.

### Colocalization analysis

We wanted to evaluate the extent to which our associations might represent single causal variants across multiple traits and used conditional association at the locus to evaluate this. For each variant within 500 kb of our lead SNPs in at least one metabolite, we ran a conditional analysis for the variants within 1 Mb of the gene body of our putative target gene. Then we ran the following association test in plink2:

`plink2 --glm cols=chrom,pos,ref,alt,a1freq,firth,test, nobs,orbeta,se,ci,tz,p hide-covar omit-ref`

`--pfile <imputed genotypes>`

`--covar <age/sex/PCs>`

`--keep <94464 European-ancestry individuals in the BOLT-LMM GWAS>`

`--out conditional/$gene/$snp`

`--pheno <technical-residualized traits>`

`--extract <(variants within 1Mb of gene body)`

`--condition <CONDITIONAL SNP>`

For single SNP conditioning tests and `--condition-list` for conditioning on multiple variants. Associations were visually inspected to detect highly linked variants and conditioning tests were repeated with top associations in any of the key traits until there were no significant variants remaining.

For the *PCCB* vignette, additional traits were included in the analysis, including fatty acids and lipids in the Nightingale-assayed individuals and clinical biomarkers in the full cohort of European-ancestry UK Biobank participants, where traits were residualized as previously described [7]. We further included a GWAS for “hard” CAD as previously defined [60], for which results were qualitatively similar when evaluating “soft” CAD (including angina cases) and employing only EHR-based diagnoses (rather than additionally including self reported case status). Results for “hard” CAD are shown in the supplement.

### Pathway diagrams

Diagrams were drawn using Affinity Design, and molecular structures were made using ChemDraw. Pathway information was curated from GO [45, 46], KEGG [47], or Reactome MSigDB [48, 49], and Stanford’s Human Metabolism Map [54], along with manual curation from public domain biochemistry knowledge (Supplementary Table S2).

## Acknowledgements

We thank Alyssa Lyn Fortier, Hanna M. Ollila, Shoa Clarke and other members of the Pritchard lab and Assimes lab for helpful discussions. The authors are grateful to UK Biobank and its participants for access to data to undertake this study (Project #30418 and #24983). Nightingale Health Plc is acknowledged for early access to the UK Biobank NMR biomarker data. C.J.S. is supported by a National Science Foundation Graduate Research Fellowship and Stanford’s Knight-Hennessy Scholars Program. This work was supported by NIH grants 5R01HG011432 and 5R01AG066490 (to J.K.P.).

## Data availability

GWAS summary statistics generated for this study will be deposited on GWAS Catalog. There will also be an interactive website app made available for interested readers to project the GWAS results for these 16 metabolites for their variant of interest onto the pathway diagram in Figure 1.

## Supplementary Figures

**Figure S1:**
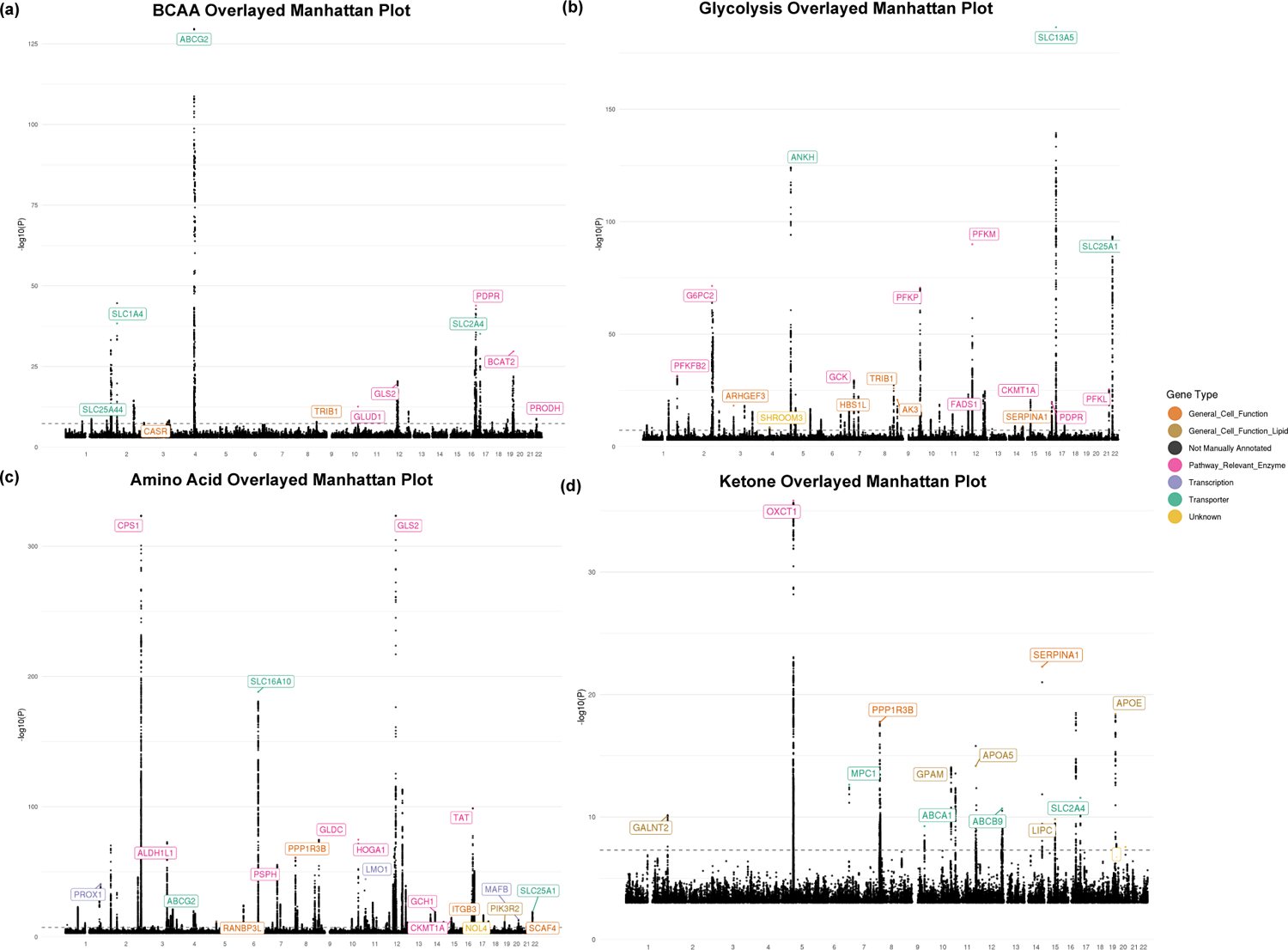
Manhattan plots. Manhattan plots for each metabolite overlayed by biochemical group. Coloring reflects gene type assignment for where relevant.

**Figure S2:**
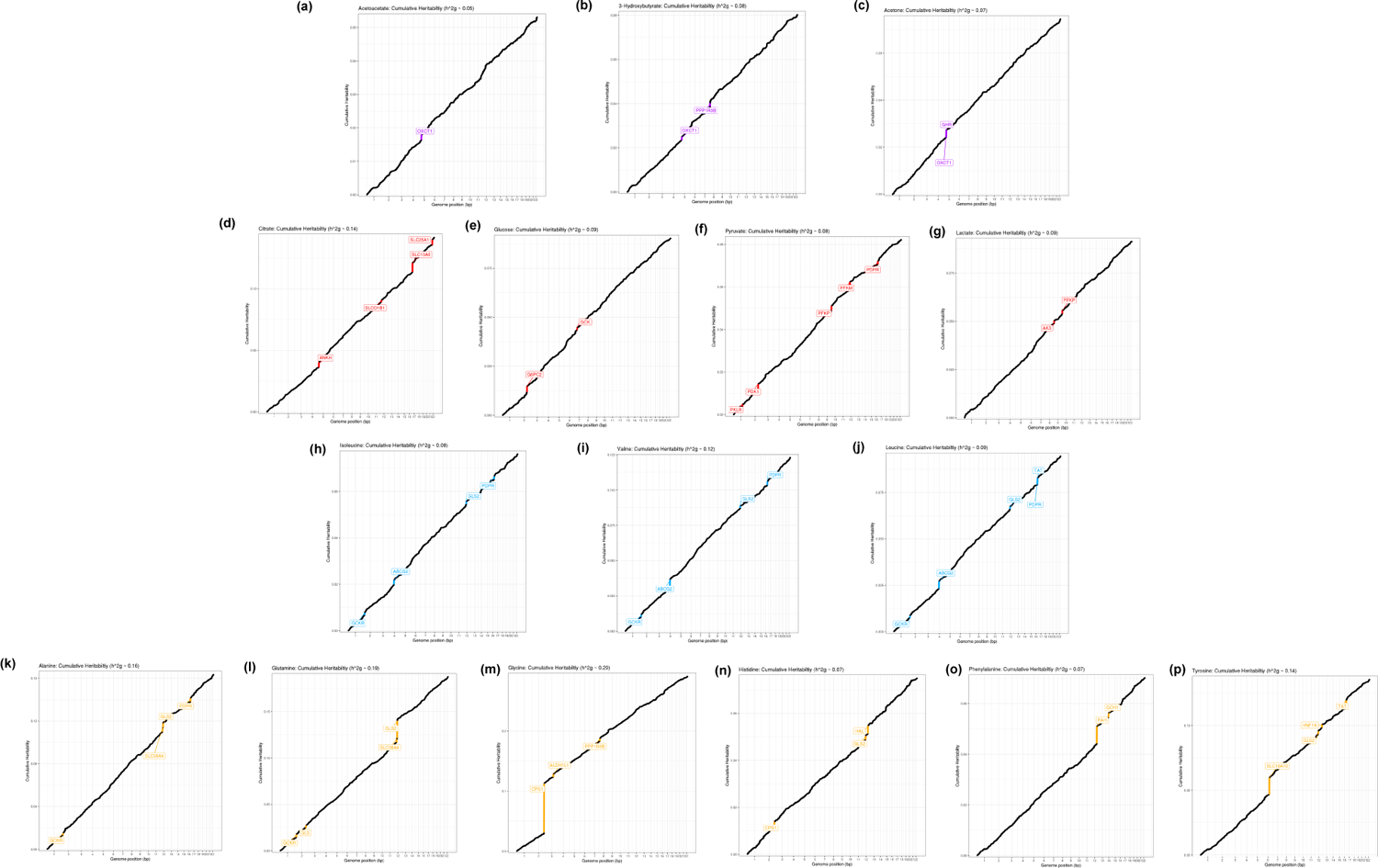
HESS heritability plots. Cummulative HESS heritability plots for the 16 metabolites, colored by biochemical group.

**Figure S3:**
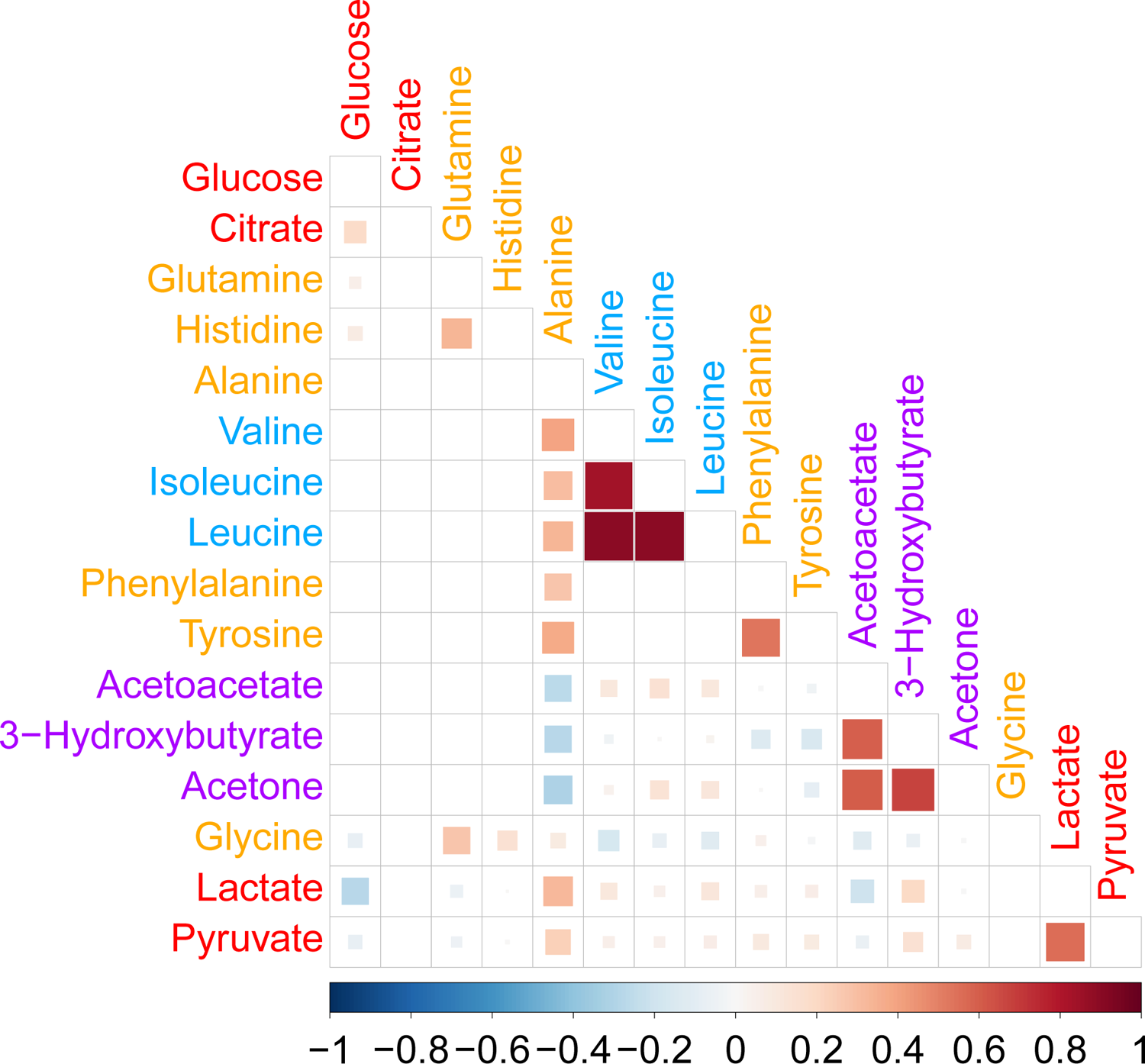
Phenotypic correlation. Correlation matrix of the residualized phenotype levels for the 16 metabolites. Metabolites are colored by biochemical group.

**Figure S4:**
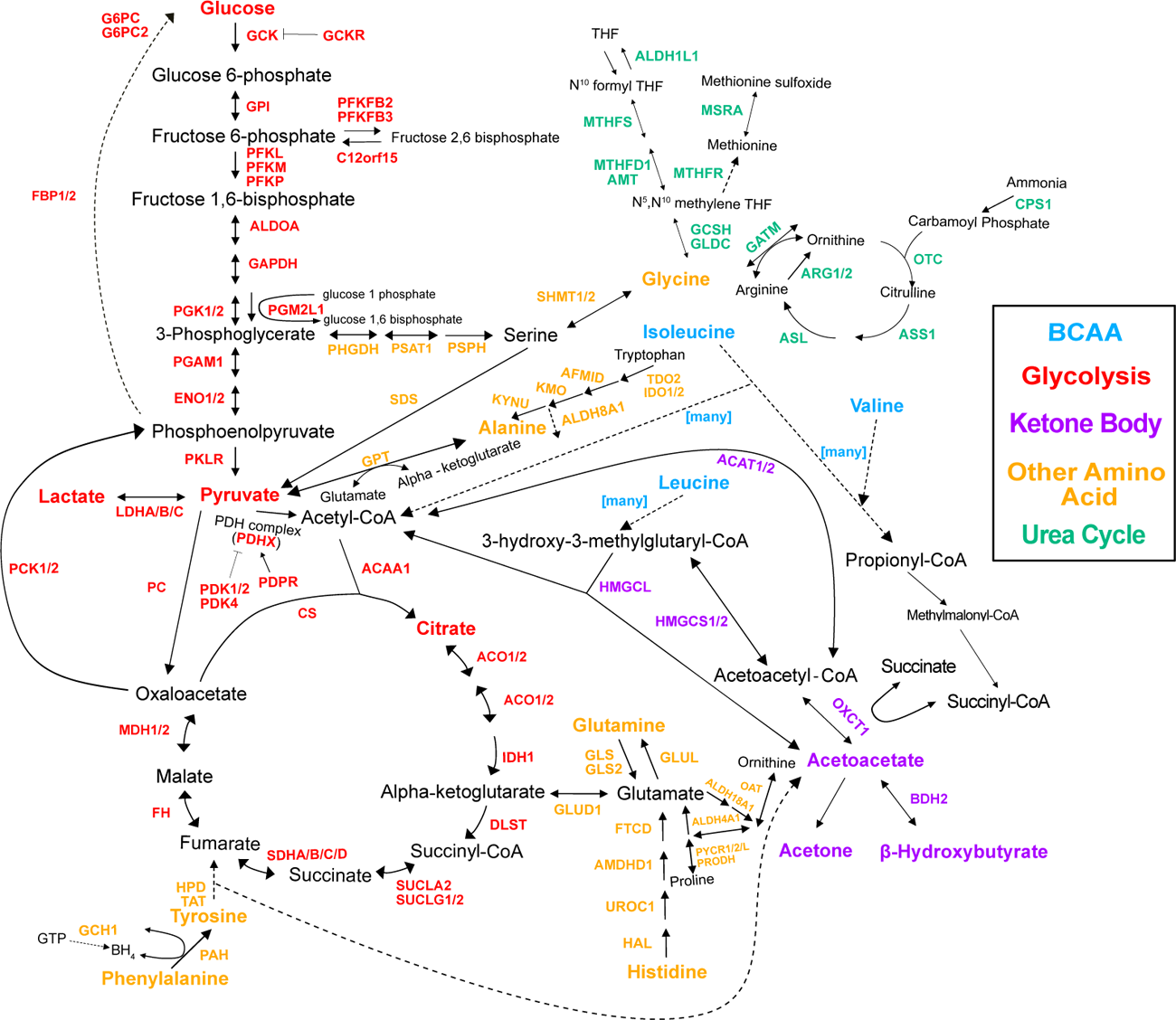
Genes in the pathway. Pathway diagram showing the genes in the pathway, regardless of whether there was a variant annotated with them or not.

**Figure S5:**
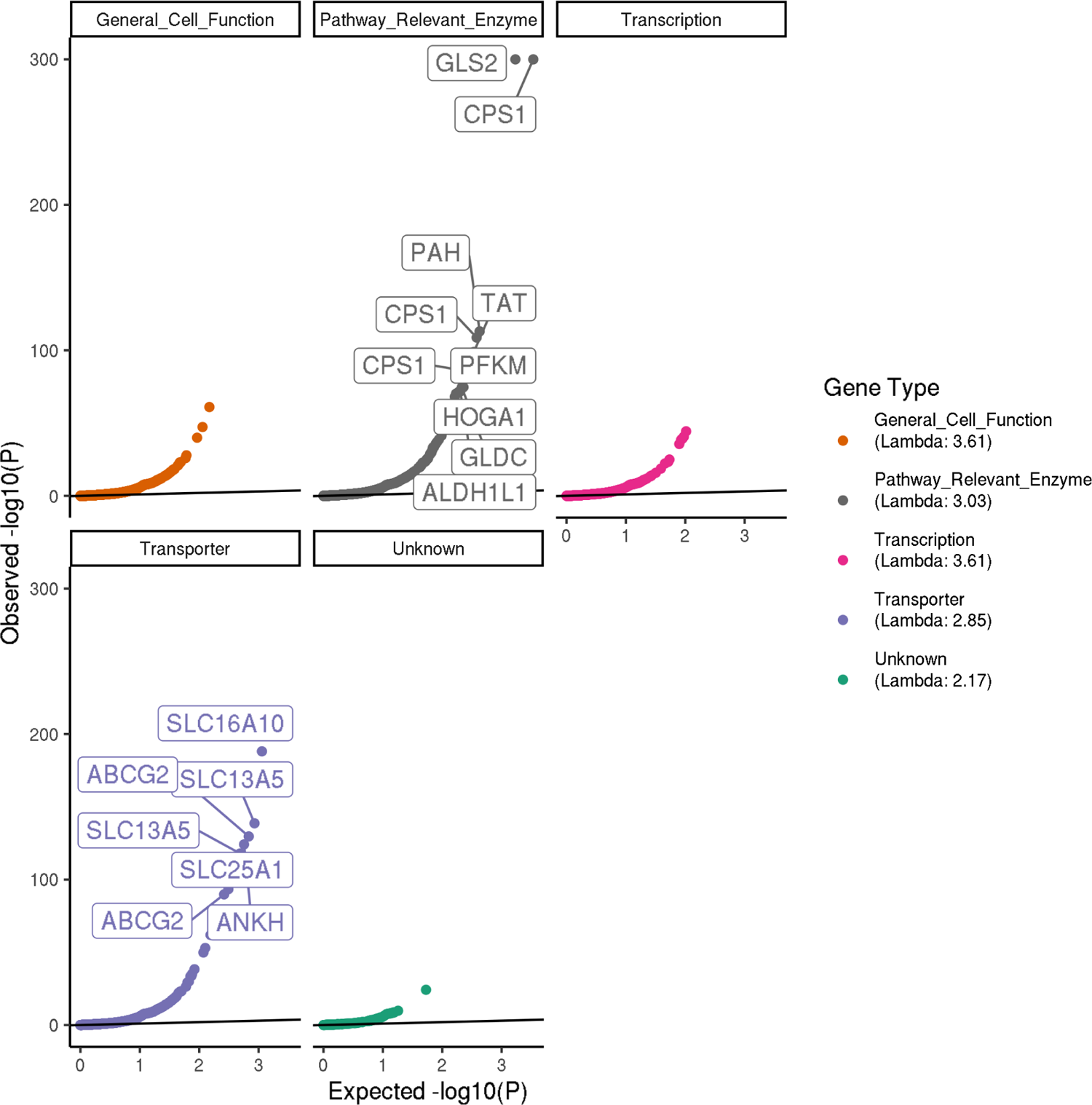
Quantile-quantile plots. Quantile-quantile plots showing the observed -log10(P-value) from the 16 GWAS in each metabolite for the 213 Metabolite GWAS Hits. The most significant trait for each SNP is excluded, and the plots are faceted by gene type. Note that for plotting purposes, P-values were set to a minimum of 1e-300. Variants with quantile < 0.005 were labeled with their gene annotation.

**Figure S6:**
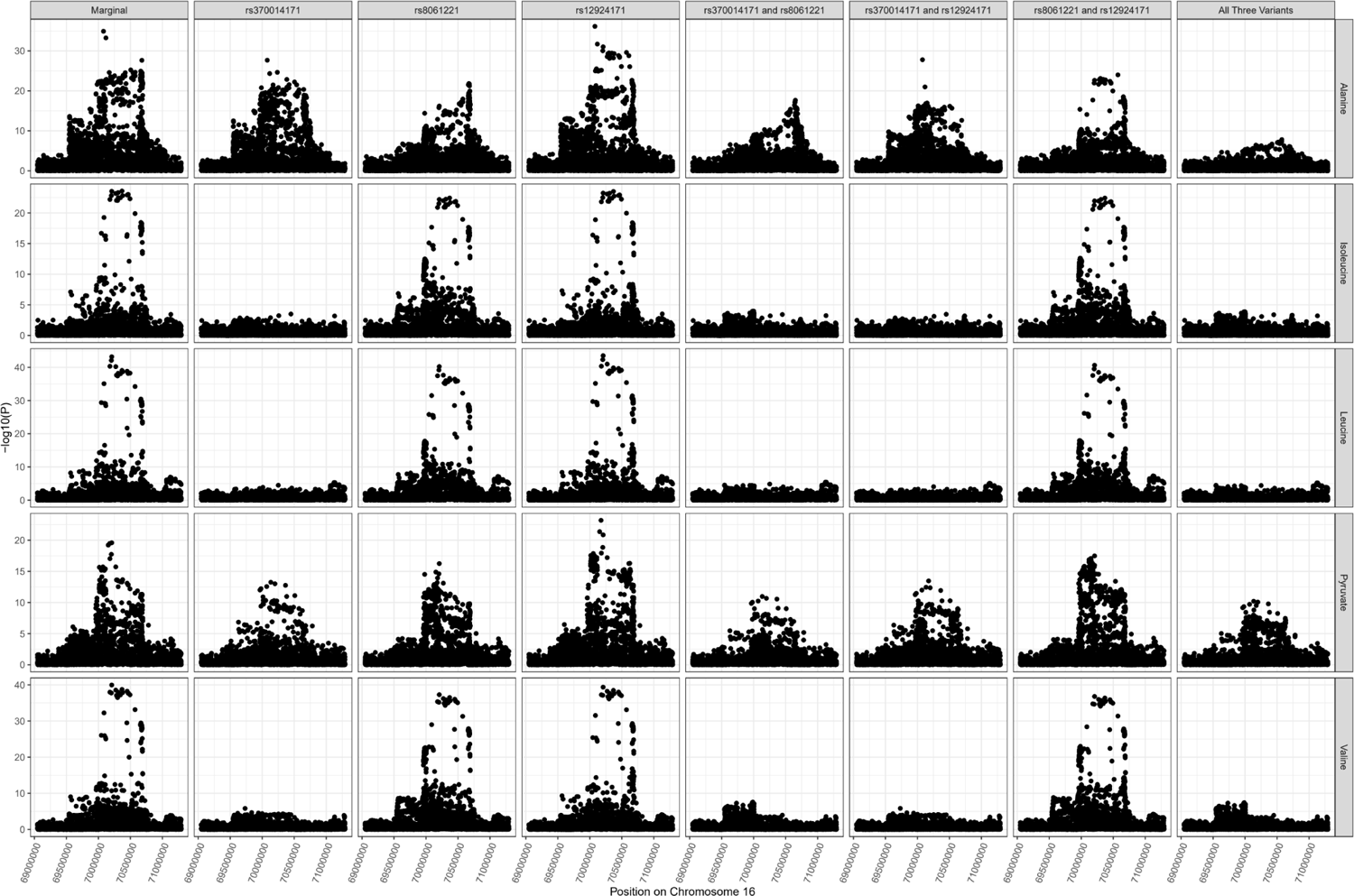
PDPR locus colocalization. Colocalization results for PDPR locus demonstrating pleiotropic effects of variant rs370014171 on alanine (one of multiple independent associations at this locus), pyruvate, isoleucine, valine, and leucine.

**Figure S7:**
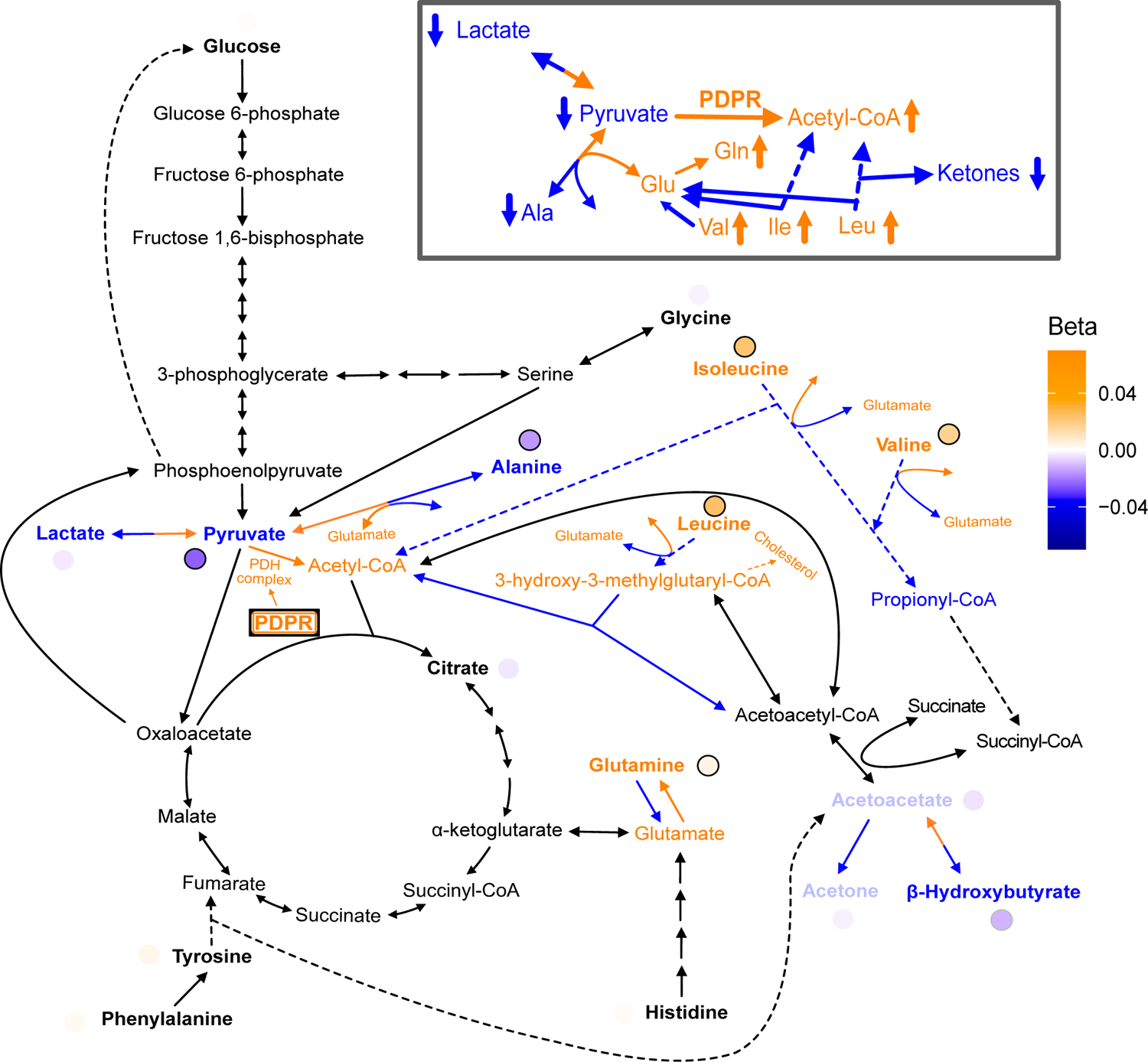
Full Pathway Results for PDPR. Full pathway results and possible mechanism for discordant variant rs370014171 with gene annotation PDPR. rs370014171 (PDPR): The discordant variant rs370014171 had gene annotation *PDPR*, which encodes the regulatory unit of the protein pyruvate dehydrogenase phosphatase which is responsible for the activation of the pyruvate dehydrogenase (PDH) complex that catalyzes the conversion of pyruvate to acetyl-CoA (Supplementary Expanded Pathway Figure S7). One possible mechanism for this variant is that it is increasing PDPR activity leading to increased conversion of pyruvate to acetyl-CoA, resulting in decreased pyruvate levels and, by compensation, decreased lactate levels. In addition, to compensate for the decreased pyruvate levels, there could be an increased conversion of alanine to pyruvate and glutamate. This would cause a decrease in levels of alanine and an increase in levels of glutamate. In response to the increased acetyl-CoA, there could be decreased breakdown of metabolites that are normally broken down for its production, including isoleucine and leucine, resulting in an increase in their levels. With the increase in acetyl-CoA levels, less HMG-CoA is broken down to acetyl-CoA. The reaction of HMG-CoA to acetyl-CoA also produces acetoacetate, so this decrease results in a decrease in acetoacetate, as well as the other ketone bodies (acetone and *β*-hydroxybutyrate), which are directly downstream of acetoacetate. Some of the increase in HMG-CoA (from its decreased breakdown) leads to an increase in cholesterol. Meanwhile, alanine to glutamate is an important regulator of BCAA levels since the first step in BCAA breakdown is a reversible conversion to glutamate. An increase in glutamate’s levels means again less BCAA will need to be broken down, which is another potential reason for increased levels of the BCAAs (isoleucine, leucine, and valine).

**Figure S8:**
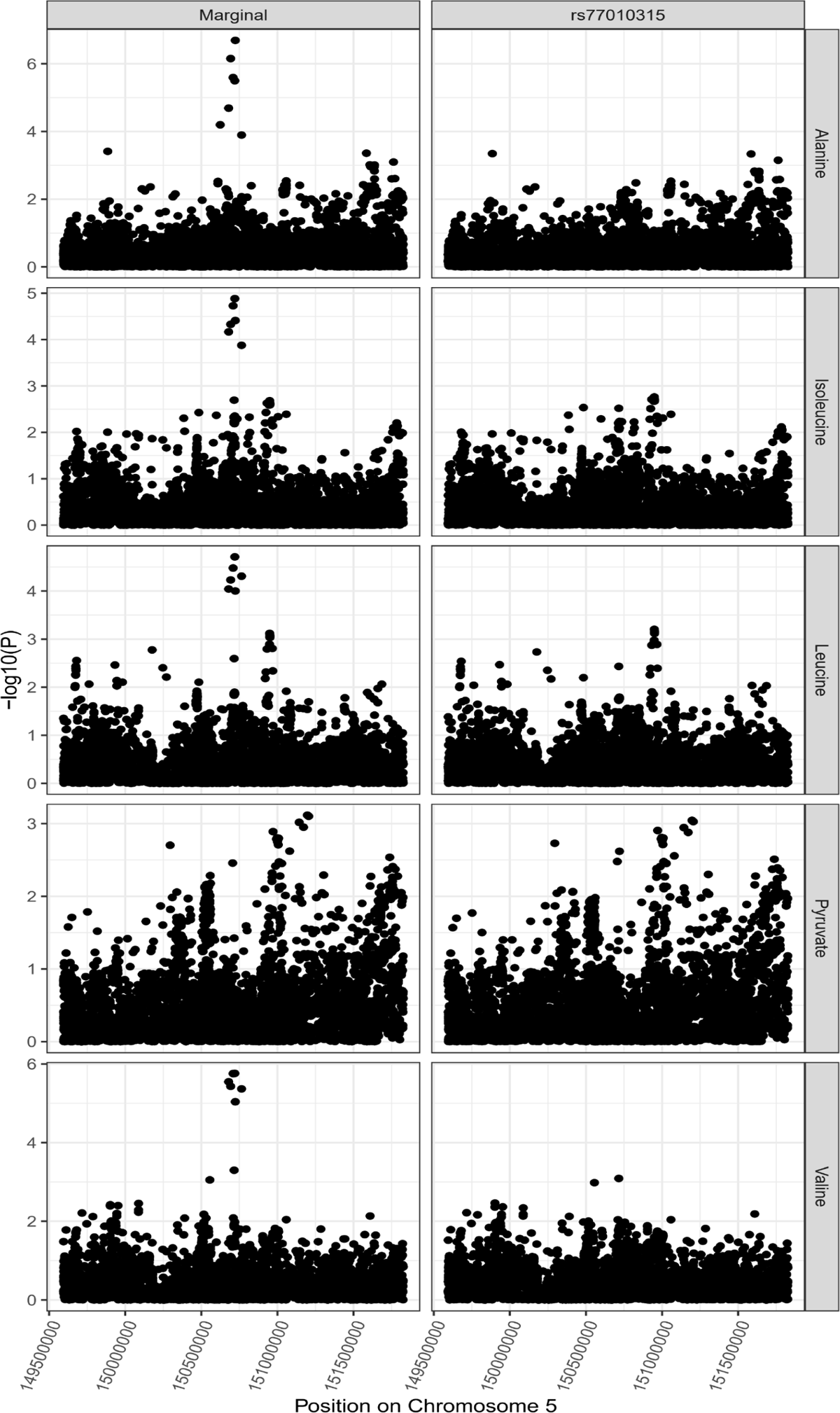
SLC36A2 locus colocalization. Colocalization results for SLC36A2 locus demonstrating pleiotropic effects of variant rs77010315 on alanine, pyruvate, isoleucine, valine and leucine.

**Figure S9:**
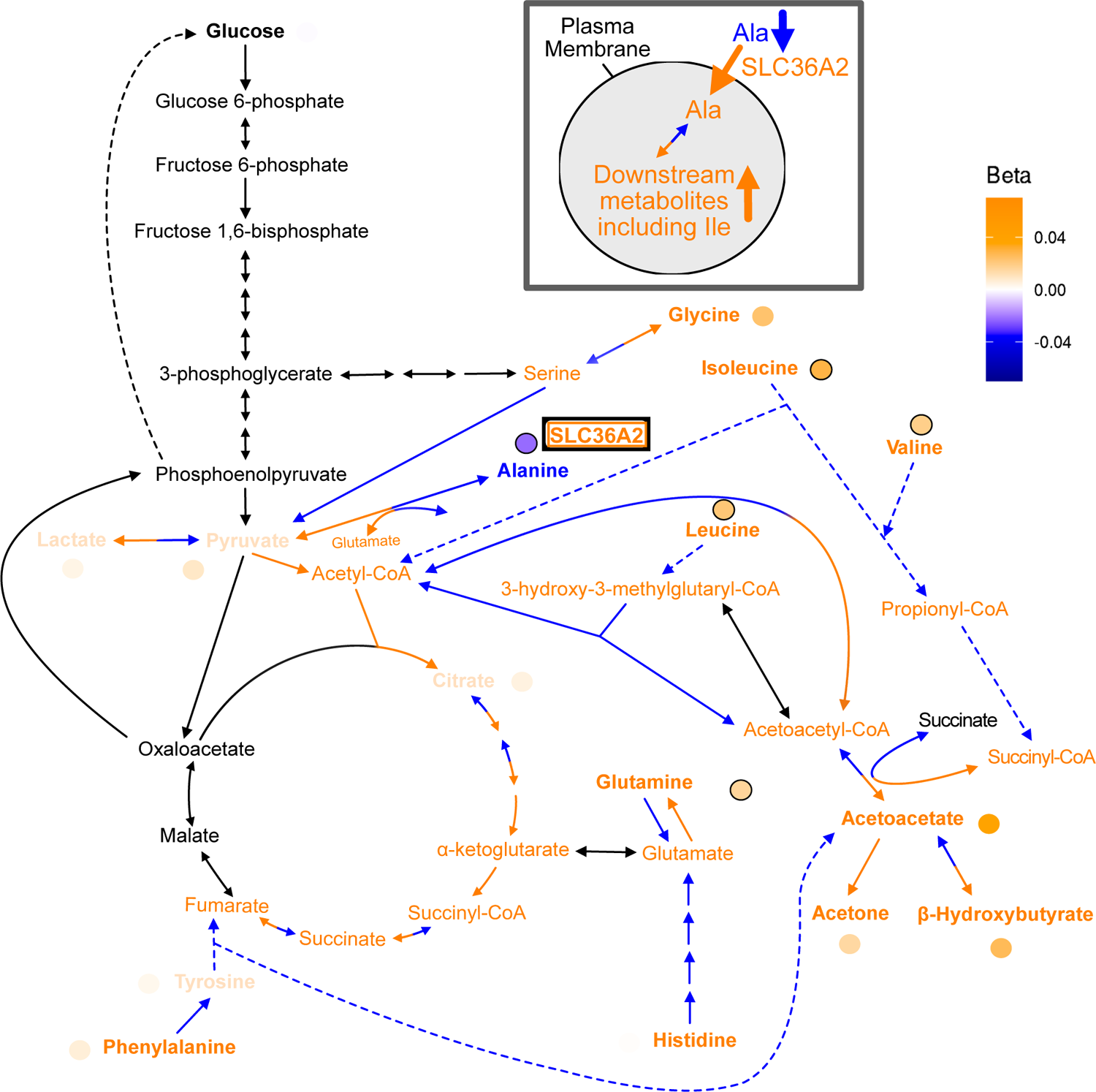
Full Pathway Results for SLC36A2. Full pathway results and possible mechanism for discordant variant rs77010315 with gene annotation SLC36A2. rs77010315 (SLC36A2): The discordant variant rs77010315 is a missense variant in *SLC36A2*, which encodes a transporter for small amino acids such as alanine (Supplementary Expanded Pathway Figure S9). We hypothesize this variant is discordant because it increases the activity of SLC36A2, leading to increased transport of alanine into cells. This results in a decrease in levels of alanine in the blood, but results in increased intracellular conversion of alanine to pyruvate and glutamate. Pyruvate can then be reversibly converted to lactate, resulting in an increase in lactate levels, as well as converted to acetyl-CoA. The increased pyruvate results in an increase in glycine levels because less needs to be broken down to produce it. The increase in acetyl-CoA leads to an increase in its downstream citric acid cycle intermediates including citrate, fumarate, succinyl-CoA and alpha-ketoglutarate. The increased glutamate leads to an increase in glutamine and the increase in fumarate leads to an increase in tyrosine and phenylalanine. The increase in acetyl-CoA also leads to an increase in isoleucine, HMG-CoA and leucine since less needs to be broken down to produce it, and to increased acetoacetyl-CoA since more acetyl-CoA is available to be converted to it. Acetoacetyl-CoA can then be converted to acetoacetate leading to an increase in all three ketone bodies (acetoacetate, acetone and *β*-hydroxybutyrate) as well as succinyl-CoA. Finally, the increased succinyl-CoA leads to an increase in valine because less valine needs to be broken down to produce it.

**Figure S10:**
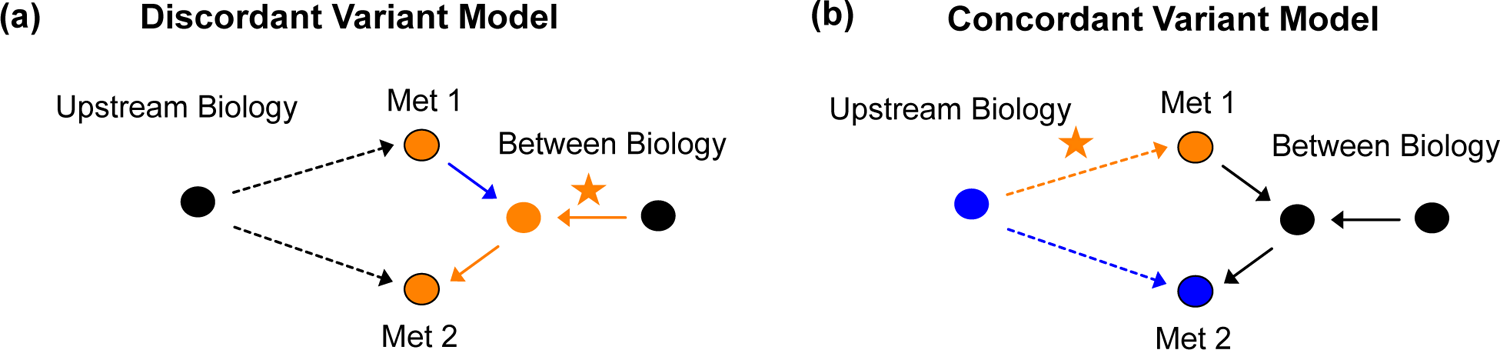
Negative Genetic Correlation Variant Models. Model for an example discordant **a.** and an example concordant **b.** variant when there is an overall negative genetic correlation.

**Figure S11:**
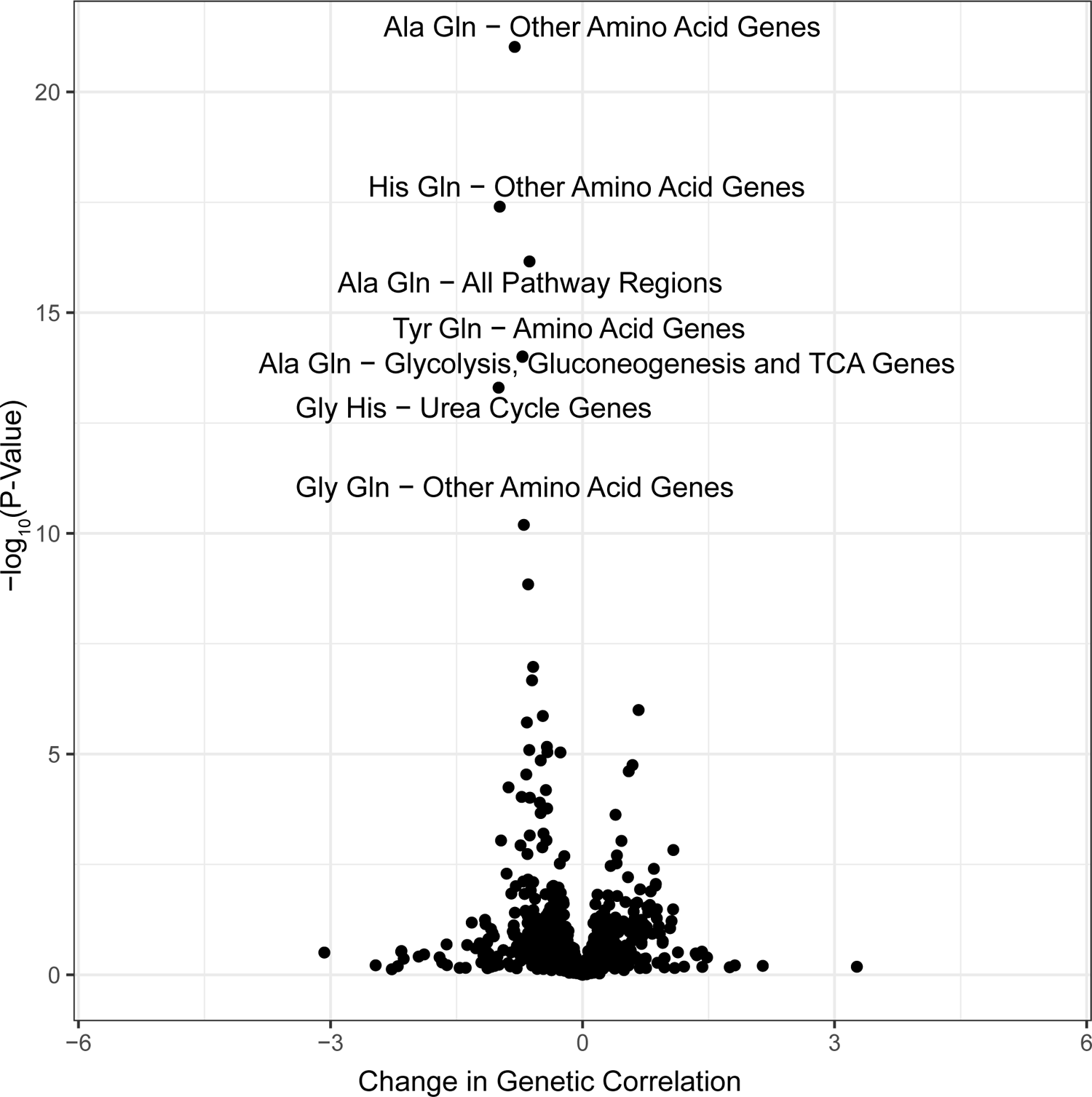
Genetic correlation results. P-values for all metabolite pairs for all pathway regions for Haseman-Elston regression genetic correlation analysis.

**Figure S12:**
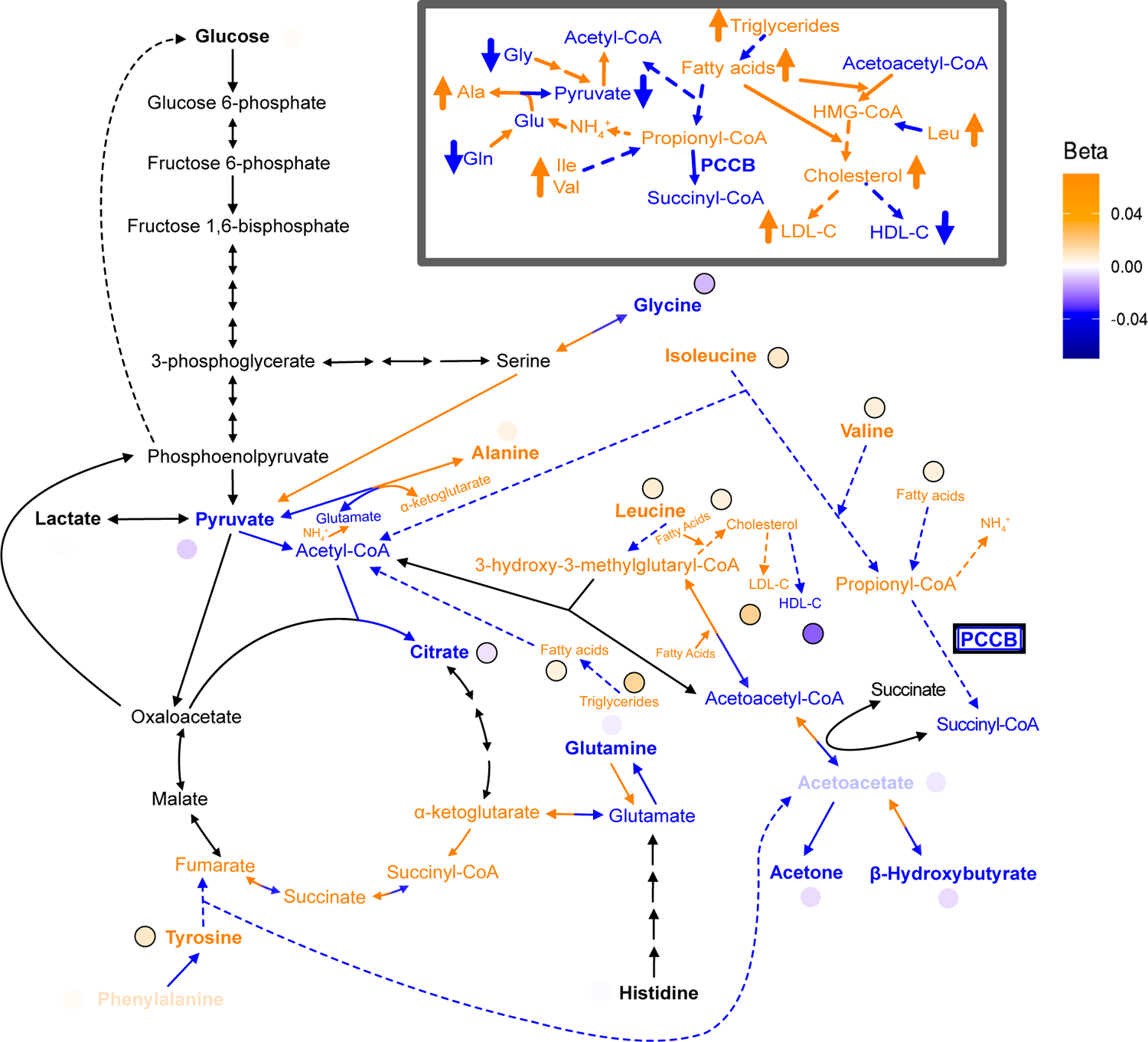
Full Pathway Results for PCCB. Full pathway results and possible mechanism for disease-associated variant rs61791721 with gene annotation PCCB. rs61791721 (PCCB): The variant rs61791721 has gene assignment *PCCB*, which encodes a protein that catalyzes the conversion of propionyl-CoA to succinyl-CoA (Supplementary Expanded Pathway Figure S12). The hypothesized mechanism is that the variant decreases PCCB activity, resulting in lower levels of succinyl-CoA and increased propionyl-CoA. The increased propionyl-CoA results in excess ammonium being produced because propionyl-CoA inhibits N-acetylglutamate synthase, which is an important cofactor for the enzyme (carbamoyl phosphate synthetase) that catalyzes the first step in the urea cycle for ammonium capture. Alanine is a reservoir for nitrogen waste so there is an increase in pyruvate and glutamate to alanine and alpha-ketoglutarate to capture the toxic ammonium [34, 35]. More glycine is then broken down in response to the decrease in pyruvate levels and more glutamine is converted to glutamate, leading to a decrease in glycine and glutamine levels. Conversely, the increased levels of propionyl-CoA mean less valine and isoleucine need to be broken down, resulting in an increase in their levels. In addition, the increased alpha-ketoglutarate increases downstream citric acid cycle intermediates meaning less tyrosine needs to be broken down to produce fumarate. Also in response to increased propionyl-CoA, fewer fatty acids are broken down, resulting in decreased acetyl-CoA, which is typically the downstream product, decreased citrate, which is one step downstream of acetyl-CoA, increased total fatty acid levels and increased total triglyceride levels. This increase may stimulate the activity of HMG-CoA reductase and synthase, resulting in increased conversion of ketone bodies to acetoacetyl-CoA to HMG-CoA and cholesterol. This results in decreased levels of ketone bodies and increased HMG-CoA and cholesterol. Increased HMG-CoA leads to increased leucine, while increased cholesterol leads to an increase in LDL-C and a decrease in HDL-C. While it is not necessarily intuitive that an increase in cholesterol levels would decrease HDL-C, it is possible that this increase activates the Rho A signal transduction pathway and suppresses peroisome proliferator-activated receptor alpha which then decreases the amount of ApoA1, as the reverse may explain why patients on statins may experience an increase in HDL-C despite a decrease in total cholesterol [61]. ApoA1 is an essential protein for HDL and thus a decrease in its levels would result in a decrease in HDL-C. We also found that *PCCB* was the strongest eGene in GTEx for rs61791721, with an effect shared across most tissues (Supplementary Figure S15). Note, we additionally ran GWAS for total triglycerides, total fatty acids, HDL cholesterol, and LDL cholesterol as part of the interpretation of the *PCCB* variant because they are important metabolites in the discussion of cardiometabolic disease (Supplementary Figure S16).

**Figure S13:**
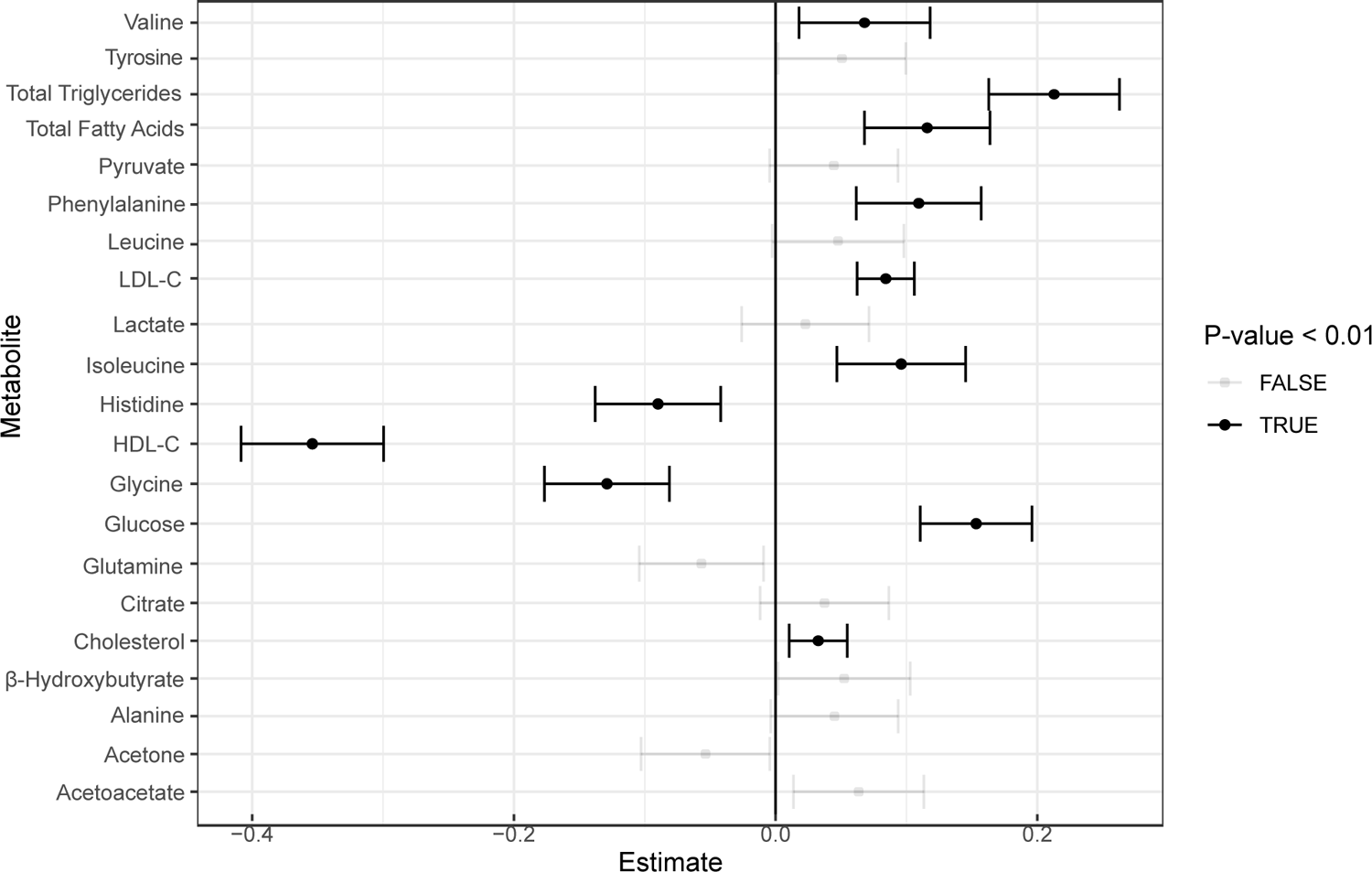
Incident analysis. Forest plot for CAD incident analysis for the 16 metabolites as well as HDL-C, LDL-C, total triglycerides, cholesterol, and total fatty acids in the UK Biobank.

**Figure S14:**
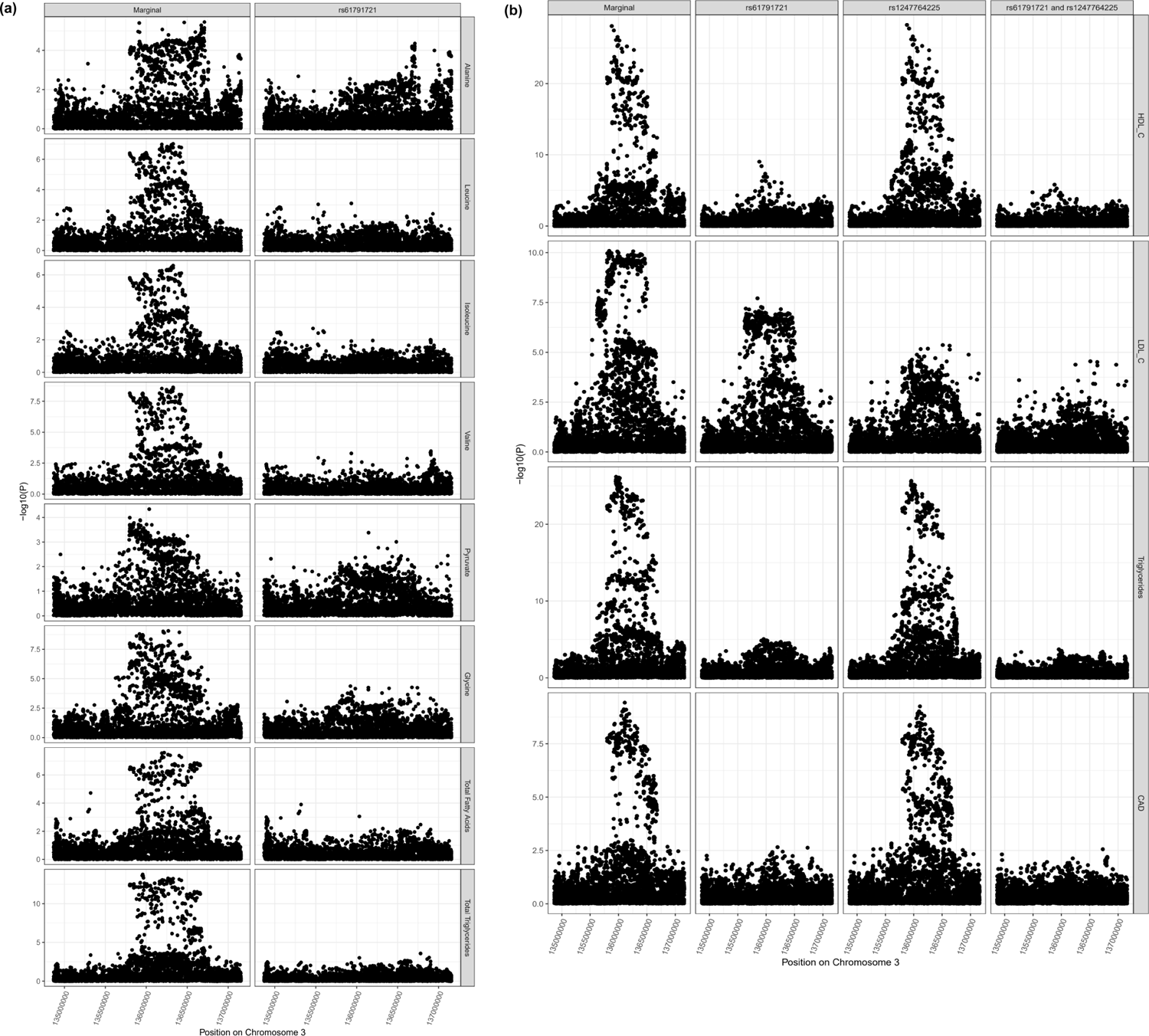
PCCB locus colocalization. Colocalization results for PCCB locus demonstrating pleiotropic effects of variant rs61791721 on alanine, glycine, pyruvate, isoleucine, valine, leucine, total fatty acids, total triglycerides, total free cholesterol, the UK Biobank clinical measures of HDL-C, LDL-C and triglycerides, and CAD.

**Figure S15:**
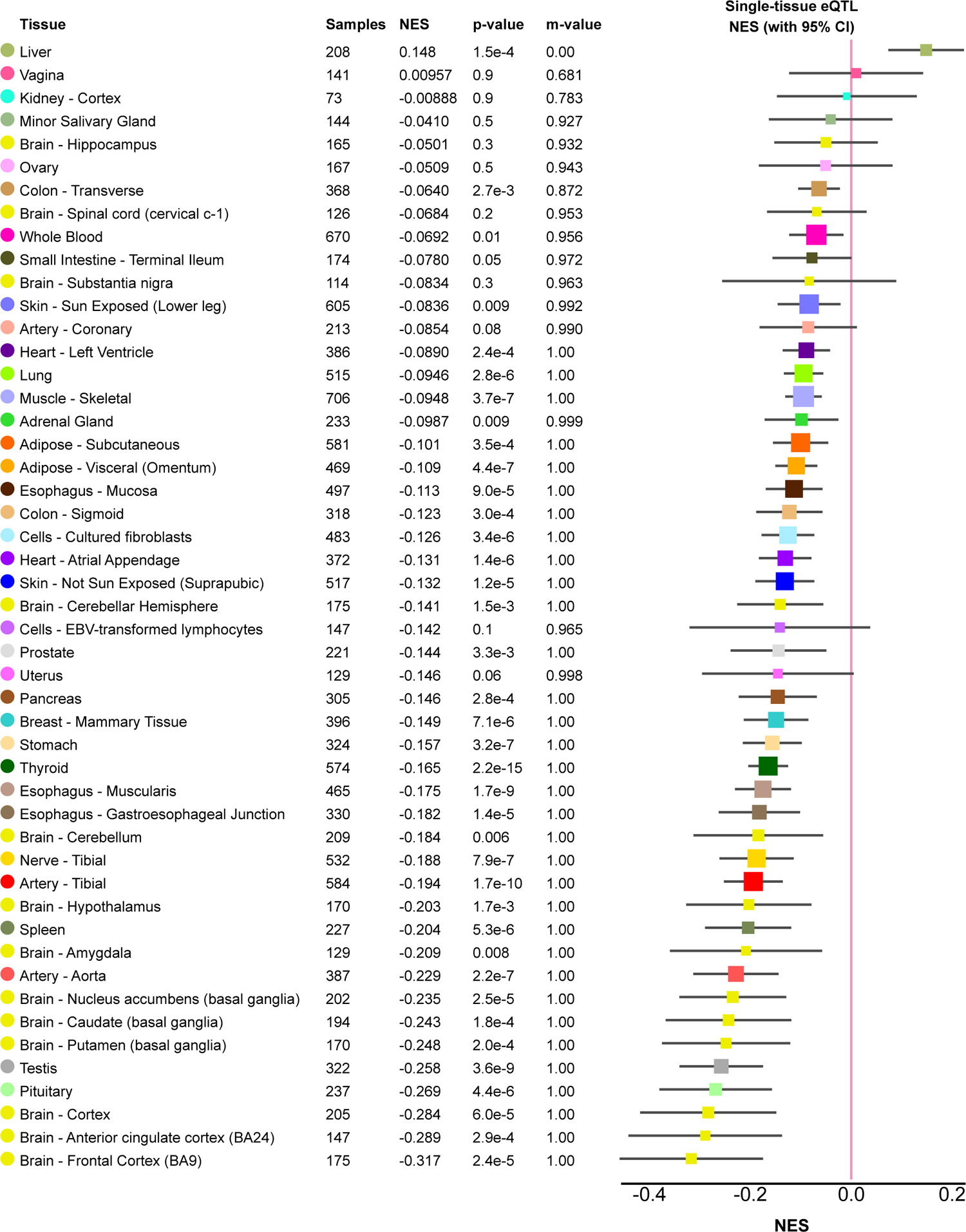
rs61791721 eQTL effects. Tissue eQTL effects for the PCCB variant (rs61791721) in GTEx.

**Figure S16:**
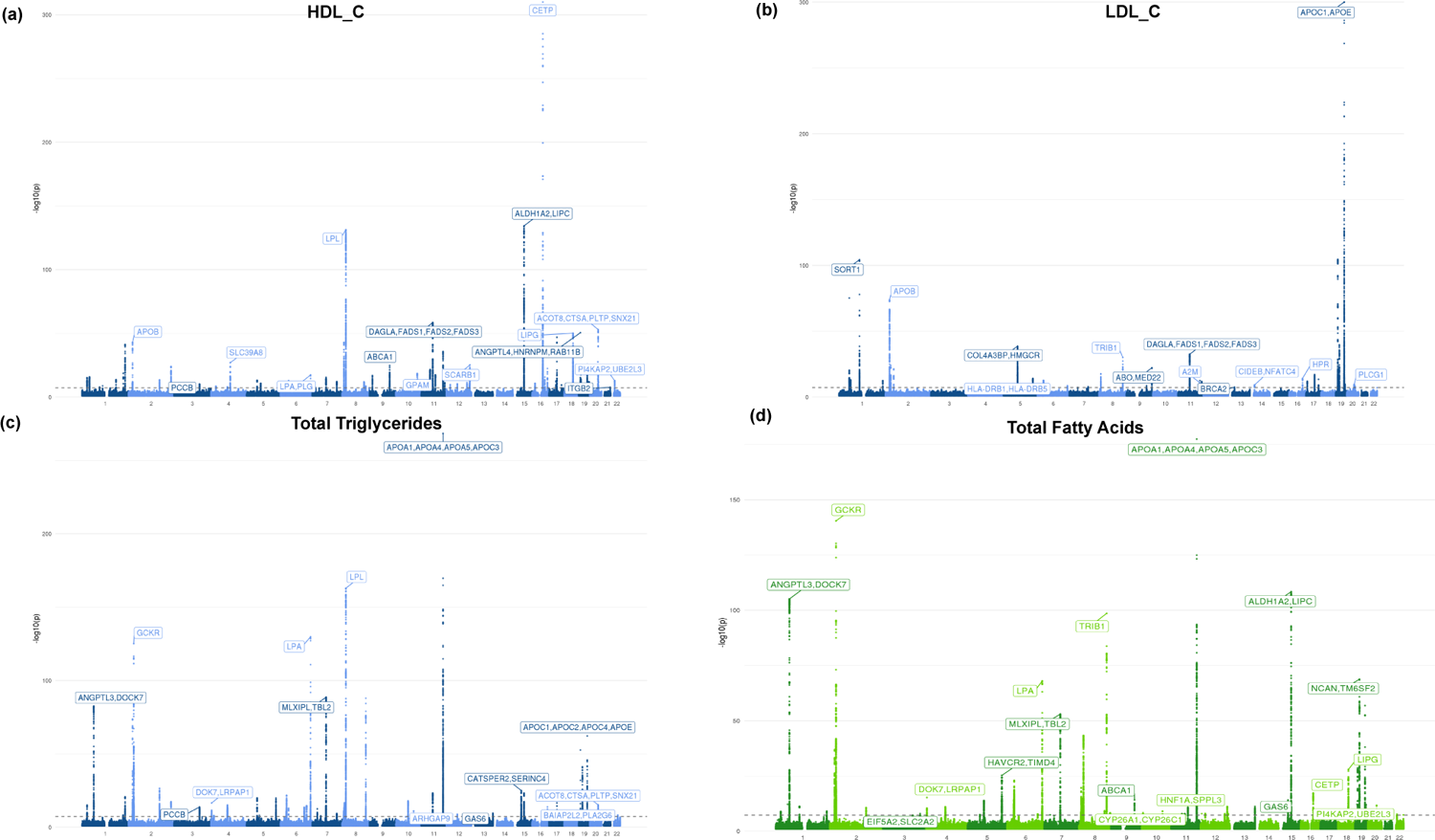
Additional manhattan plots. Manhattan plots for (a) HDL_C, (b) LDL_C, (c) total triglycerides, and (d) total fatty acids. Blue coloring represents metabolites belonging to the lipid group and green belongs to fatty acid group.

## Supplementary Tables

**Table S1: Metabolite HESS heritabilities.** Heritibility results from HESS for each metabolite. (File: hess_scaled_heritabilities.tsv)

**Table S2: Gene function sources.** Sources for biochemical characterization of genes mentioned in the variant vignettes. (File: gene_biochemistry_sources.xlsx)

**Table S3: Metabolite GWAS hits annotation.** Annotation for the 213 metabolite GWAS hits, including the assigned gene, assigned gene type, variant classification, and nearest gene. (File: rigidmetabolites_sig_pruned_snplist_manual_annotation_closestgene_disttss.tsv)

**Table S4: Gene biochemical groups.** The number of significant (P < 1e-4) metabolite associations each metabolite GWAS gene had for each biochemical group. For a given gene, only biochemical groups that had at least one significant metabolite association were listed. (File: gene_metgroup_assignments_all.tsv)

**Table S5: Ancestry-inclusive GWAS hits.** List of additional metabolite GWAS hits from the ancestry-inclusive analysis that were not present in the European-only GWAS results. (File: newsnps_annotated_rigid_novel_withsumstats.tsv)

**Table S6: Discordant variant annotation.** List of each discordant variant-metabolite association, including the variant annotations and relevant metabolite pair genetic correlation and GWAS summary statistics. (File: disvar_annotation.tsv)

**Table S7: Local genetic correlation results.** Combined results for different methods of calculating the local genetic correlation for different pathways for alanine and glutamine, demonstrating the consistency across the different approaches. (File: combined_localgencor_methods.tsv)

**Table S8: Metabolite associations with CAD.** Literature evidence and citations for metabolite associations with CAD. (File: met_to_disease.xlsx)

**Table S9: PCCB GWAS results.** GWAS summary statistics for rs61791721 (PCCB) in the 16 metabolites. (File: PCCB_sumstats.tsv)

## References

1. Nadia Solovieff, Chris Cotsapas, Phil H. Lee, Shaun M. Purcell, and Jordan W. Smoller. Pleiotropy in complex traits: challenges and strategies. Nat Rev Genet, 14(7):483–495. 2013-07.

2. Brendan K. Bulik-Sullivan, Po-Ru Loh, Hilary K. Finucane, Stephan Ripke, Jian Yang, Nick Patterson, Mark J. Daly, Alkes L. Price, and Benjamin M. Neale. LD score regression distinguishes confounding from polygenicity in genome-wide association studies. Nat Genet, 47(3):291–295. 2015-03.

3. Huwenbo Shi, Nicholas Mancuso, Sarah Spendlove, and Bogdan Pasaniuc. Local genetic correlation gives insights into the shared genetic architecture of complex traits. The American Journal of Human Genetics, 101(5):737–751. 2017-11-02.

4. Curtis R. Warren, John F. O’Sullivan, Max Friesen, et al. Induced pluripotent stem cell differentiation enables functional validation of GWAS variants in metabolic disease. Cell Stem Cell, 20(4):547–557.e7. 2017-04.

5. Nasa Sinnott-Armstrong, Isabel S. Sousa, Samantha Laber, et al. A regulatory variant at 3q21.1 confers an increased pleiotropic risk for hyperglycemia and altered bone mineral density. Cell Metab, 33(3):615–628.e13. 2021-03-02.

6. Christian Gieger, Ludwig Geistlinger, Elisabeth Altmaier, et al. Genetics meets metabolomics: A genome-wide association study of metabolite profiles in human serum. PLOS Genetics, 4(11):e1000282. 2008-11-28.

7. Nasa Sinnott-Armstrong, Sahin Naqvi, Manuel Rivas, and Jonathan K Pritchard. GWAS of three molecular traits highlights core genes and pathways alongside a highly polygenic background. eLife, 10:e58615. 2021-02-15.

8. Mathias Woidy, Ania C. Muntau, and Søren W. Gersting. Inborn errors of metabolism and the human interactome: a systems medicine approach. J Inherit Metab Dis, 41(3):285–296. 2018-05-01.

9. So-Youn Shin, Eric B. Fauman, Ann-Kristin Petersen, et al. An atlas of genetic influences on human blood metabolites. Nat Genet, 46(6):543–550. 2014-06.

10. Victoria Au Yeung. Common ‘inborn errors’ of metabolism in the general population. University of Cambridge. Defended 2021-03-23.

11. Heli Julkunen, Anna Cichońska, P Eline Slagboom, Peter Würtz, and Nightingale Health UK Biobank Initiative. Metabolic biomarker profiling for identification of susceptibility to severe pneumonia and COVID-19 in the general population. eLife, 10:e63033. 2021-05-04.

12. Lori Laffel. Ketone bodies: a review of physiology, pathophysiology and application of monitoring to diabetes. Diabetes/Metabolism Research and Reviews, 15(6):412–426. 1999.

13. Marta Guasch-Ferré, José L Santos, Miguel A Martínez-González, et al. Glycolysis/gluconeogenesis- and tricarboxylic acid cycle–related metabolites, mediterranean diet, and type 2 diabetes. The American Journal of Clinical Nutrition, 111(4):835–844. 2020-04-01.

14. P. Newsholme, K. Bender, A. Kiely, and L. Brennan. Amino acid metabolism, insulin secretion and diabetes. Biochemical Society Transactions, 35(5):1180–1186. 2007-10-25.

15. Aldons J. Lusis and James N. Weiss. Cardiovascular networks. Circulation, 121(1):157–170. 2010-01-05.

16. Matthew J Watt, Paula M Miotto, William De Nardo, and Magdalene K Montgomery. The liver as an endocrine organ—linking NAFLD and insulin resistance. Endocrine Reviews, 40(5):1367– 1393. 2019-10-01.

17. Rozenn N. Lemaitre, Toshiko Tanaka, Weihong Tang, et al. Genetic loci associated with plasma phospholipid n-3 fatty acids: A meta-analysis of genome-wide association studies from the CHARGE consortium. PLOS Genetics, 7(7):e1002193. 2011-07-28.

18. Karsten Suhre, So-Youn Shin, Ann-Kristin Petersen, et al. Human metabolic individuality in biomedical and pharmaceutical research. Nature, 477(7362):54–60. 2011-09.

19. Tanya M Teslovich, Daniel Seung Kim, Xianyong Yin, et al. Identification of seven novel loci associated with amino acid levels using single-variant and gene-based tests in 8545 finnish men from the METSIM study. Human Molecular Genetics, 27(9):1664–1674. 2018-05-01.

20. Sarah E. Graham, Shoa L. Clarke, Kuan-Han H. Wu, et al. The power of genetic diversity in genome-wide association studies of lipids. Nature, pages 1–11. 2021-12-09.

21. Rico Rueedi, Roger Mallol, Johannes Raffler, David Lamparter, Nele Friedrich, Peter Vollenweider, Gérard Waeber, Gabi Kastenmüller, Zoltán Kutalik, and Sven Bergmann. Metabomatching: Using genetic association to identify metabolites in proton NMR spectroscopy. PLoS Comput Biol, 13(12):e1005839. 2017-12-01.

22. Johannes Kettunen, Ayşe Demirkan, Peter Würtz, et al. Genome-wide study for circulating metabolites identifies 62 loci and reveals novel systemic effects of LPA. Nat Commun, 7(1):11122. 2016-03-23.

23. Laura B. L. Wittemans, Luca A. Lotta, Clare Oliver-Williams, et al. Assessing the causal association of glycine with risk of cardio-metabolic diseases. Nat Commun, 10(1):1060. 2019-03-05.

24. Luca A. Lotta, Maik Pietzner, Isobel D. Stewart, et al. A cross-platform approach identifies genetic regulators of human metabolism and health. Nat Genet, 53(1):54–64. 2021-01.

25. Janne Pott, Yoon Ju Bae, Katrin Horn, et al. Genetic association study of eight steroid hormones and implications for sexual dimorphism of coronary artery disease. The Journal of Clinical Endocrinology & Metabolism, 104(11):5008–5023. 2019-11-01.

26. Anna Cichonska, Juho Rousu, Pekka Marttinen, et al. metaCCA: summary statistics-based multivariate meta-analysis of genome-wide association studies using canonical correlation analysis. Bioinformatics, 32(13):1981–1989. 2016-07-01.

27. Sanni E. Ruotsalainen, Juulia J. Partanen, Anna Cichonska, et al. An expanded analysis framework for multivariate GWAS connects inflammatory biomarkers to functional variants and disease. Eur J Hum Genet, 29(2):309–324. 2021-02.

28. Guanghao Qi and Nilanjan Chatterjee. Heritability informed power optimization (HIPO) leads to enhanced detection of genetic associations across multiple traits. PLOS Genetics, 14(10):e1007549. 2018-10-05.

29. Po-Ru Loh, George Tucker, Brendan K Bulik-Sullivan, et al. Efficient bayesian mixed-model analysis increases association power in large cohorts. Nat Genet, 47(3):284–290. 2015-03.

30. Jian Yang, Beben Benyamin, Brian P. McEvoy, et al. Common SNPs explain a large proportion of the heritability for human height. Nat Genet, 42(7):565–569. 2010-07.

31. Brendan Bulik-Sullivan, Hilary K. Finucane, Verneri Anttila, et al. An atlas of genetic correlations across human diseases and traits. Nat Genet, 47(11):1236–1241. 2015-11.

32. Gleb Kichaev, Gaurav Bhatia, Po-Ru Loh, Steven Gazal, Kathryn Burch, Malika K. Freund, Armin Schoech, Bogdan Pasaniuc, and Alkes L. Price. Leveraging polygenic functional enrichment to improve GWAS power. Am J Hum Genet, 104(1):65–75. 2019-01-03.

33. Satoshi Koyama, Kaoru Ito, Chikashi Terao, et al. Population-specific and trans-ancestry genome-wide analyses identify distinct and shared genetic risk loci for coronary artery disease. Nat Genet, 52(11):1169–1177. 2020-11.

34. Parith Wongkittichote, Nicholas Ah Mew, and Kimberly A. Chapman. Propionyl-CoA carboxylase – a review. Molecular Genetics and Metabolism, 122(4):145–152. 2017-12-01.

35. L. D. Smith and U. Garg. Chapter 5 - urea cycle and other disorders of hyperammonemia. In Uttam Garg and Laurie D. Smith, editors, Biomarkers in Inborn Errors of Metabolism, pages 103–123. Elsevier.

36. Nan Wu, Lindsei K. Sarna, Sun-Young Hwang, Qingjun Zhu, Pengqi Wang, Yaw L. Siow, and Karmin O. Activation of 3-hydroxy-3-methylglutaryl coenzyme a (HMG-CoA) reductase during high fat diet feeding. Biochimica et Biophysica Acta (BBA) - Molecular Basis of Disease, 1832(10):1560–1568. 2013-10-01.

37. W H Salam, H G Wilcox, L M Cagen, and M Heimberg. Stimulation of hepatic cholesterol biosynthesis by fatty acids. effects of oleate on cytoplasmic acetoacetyl-CoA thiolase, acetoacetyl-CoA synthetase and hydroxymethylglutaryl-CoA synthase. Biochem J, 258(2):563–568. 1989-03-01.

38. Therese Tillin, Alun D. Hughes, Qin Wang, et al. Diabetes risk and amino acid profiles: cross-sectional and prospective analyses of ethnicity, amino acids and diabetes in a south asian and european cohort from the SABRE (southall and brent REvisited) study. Diabetologia, 58(5):968–979. 2015-05-01.

39. Raimo Jauhiainen, Jagadish Vangipurapu, Annamaria Laakso, Teemu Kuulasmaa, Johanna Kuusisto, and Markku Laakso. The association of 9 amino acids with cardiovascular events in finnish men in a 12-year follow-up study. J Clin Endocrinol Metab, 106(12):3448–3454. 2021-08-04.

40. Clare Bycroft, Colin Freeman, Desislava Petkova, et al. The UK biobank resource with deep phenotyping and genomic data. Nature, 562(7726):203–209. 2018-10.

41. Genevieve L. Wojcik, Mariaelisa Graff, Katherine K. Nishimura, et al. Genetic analyses of diverse populations improves discovery for complex traits. Nature, 570(7762):514–518. 2019-06.

42. Cristen J. Willer, Ellen M. Schmidt, Sebanti Sengupta, et al. Discovery and refinement of loci associated with lipid levels. Nat Genet, 45(11):1274–1283. 2013-11.

43. Céline Bellenguez, Amy Strange, Colin Freeman, Peter Donnelly, and Chris C.A. Spencer. A robust clustering algorithm for identifying problematic samples in genome-wide association studies. Bioinformatics, 28(1):134–135. 2012-01-01.

44. Christopher C Chang, Carson C Chow, Laurent CAM Tellier, Shashaank Vattikuti, Shaun M Purcell, and James J Lee. Second-generation PLINK: rising to the challenge of larger and richer datasets. GigaScience, 4(1):s13742–015–0047–8. 2015-12-01.

45. Michael Ashburner, Catherine A. Ball, Judith A. Blake, et al. Gene ontology: tool for the unification of biology. Nat Genet, 25(1):25–29. 2000-05.

46. The Gene Ontology Consortium, Seth Carbon, Eric Douglass, et al. The gene ontology resource: enriching a GOld mine. Nucleic Acids Research, 49:D325–D334. 2021-01-08.

47. Minoru Kanehisa and Susumu Goto. KEGG: Kyoto encyclopedia of genes and genomes. Nucleic Acids Res, 28(1):27–30. 2000-01-01.

48. Arthur Liberzon, Aravind Subramanian, Reid Pinchback, Helga Thorvaldsdóttir, Pablo Tamayo, and Jill P. Mesirov. Molecular signatures database (MSigDB) 3.0. Bioinformatics, 27(12):1739–1740. 2011-06-15.

49. Aravind Subramanian, Pablo Tamayo, Vamsi K. Mootha, et al. Gene set enrichment analysis: A knowledge-based approach for interpreting genome-wide expression profiles. Proc Natl Acad Sci U S A, 102(43):15545–15550. 2005-10-25.

50. Gil Stelzer, Naomi Rosen, Inbar Plaschkes, et al. The GeneCards suite: From gene data mining to disease genome sequence analyses. Current Protocols in Bioinformatics, 54(1). 2016-06.

51. Lloyd T. Elliott, Kevin Sharp, Fidel Alfaro-Almagro, Sinan Shi, Karla L. Miller, Gwenaëlle Douaud, Jonathan Marchini, and Stephen M. Smith. Genome-wide association studies of brain imaging phenotypes in UK biobank. Nature, 562(7726):210–216. 2018-10.

52. Maya Ghoussaini, Edward Mountjoy, Miguel Carmona, et al. Open targets genetics: systematic identification of trait-associated genes using large-scale genetics and functional genomics. Nucleic Acids Res, 49:D1311–D1320. 2020-10-12.

53. Edward Mountjoy, Ellen M. Schmidt, Miguel Carmona, et al. An open approach to systematically prioritize causal variants and genes at all published human GWAS trait-associated loci. Nat Genet, 53(11):1527–1533. 2021-11.

54. Pathways of human metabolism map. Stanford Med Education; https://metabolicpathways.stanford.edu/.

55. Mingcong Xu, Xuefeng Bai, Bo Ai, et al. TF-marker: a comprehensive manually curated database for transcription factors and related markers in specific cell and tissue types in human. Nucleic Acids Research, 50:D402–D412. 2022-01-07.

56. Huwenbo Shi, Gleb Kichaev, and Bogdan Pasaniuc. Contrasting the genetic architecture of 30 complex traits from summary association data. The American Journal of Human Genetics, 99(1):139–153. 2016-07-07.

57. Jack Bowden, Wesley Spiller, Fabiola Del Greco M, Nuala Sheehan, John Thompson, Cosetta Minelli, and George Davey Smith. Improving the visualization, interpretation and analysis of two-sample summary data mendelian randomization via the radial plot and radial regression. Int J Epidemiol, 47(4):1264–1278. 2018-08-01.

58. Nasa Sinnott Armstrong, Yosuke Tanigawa, David Amar, et al. Genetics of 35 blood and urine biomarkers in the UK biobank. Nat Genet, 53(2):185–194. 2021-02.

59. Matthew Stephens. False discovery rates: a new deal. Biostatistics, 18(2):275–294. 2017-04-01.

60. Michael Inouye, Gad Abraham, Christopher P. Nelson, et al. Genomic risk prediction of coronary artery disease in 480,000 adults. J Am Coll Cardiol, 72(16):1883–1893. 2018-10-16.

61. Geneviève Martin, Hélène Duez, Christophe Blanquart, Vincent Berezowski, Philippe Poulain, Jean-Charles Fruchart, Jamila Najib-Fruchart, Corine Glineur, and Bart Staels. Statin-induced inhibition of the rho-signaling pathway activates PPAR and induces HDL apoA-i. J Clin Invest, 107(11):1423–1432. 2001-06-01.

